# Species-specific oxygen sensing governs the initiation of vertebrate limb regeneration

**DOI:** 10.1101/2024.12.19.629359

**Authors:** Georgios Tsissios, Marion Leleu, Kelly Hu, Alp Eren Demirtas, Hanrong Hu, Toru Kawanishi, Evangelia Skoufa, Alessandro Valente, Antonio Herrera, Adrien Mery, Lorenzo Noseda, Haruki Ochi, Selman Sakar, Mikiko Tanaka, Fides Zenk, Can Aztekin

## Abstract

Why mammals cannot regenerate limbs, unlike amphibians, presents a longstanding puzzle in biology. We show that exposing *ex vivo* amputated embryonic mouse limbs to subatmospheric oxygen environment, or stabilizing oxygen-sensitive HIF1A enables not only rapid wound healing, but alters cellular mechanics, and reshapes the histone landscape to prime regenerative fates. Conversely, regenerative *Xenopus* tadpole limbs display low oxygen-sensing capacity, robust wound healing, a regenerative histone landscape, and glycolytic programs even under high oxygen. This reduced oxygen-sensing capacity, in stark contrast to mammals, associates with decreased HIF1A-regulating gene expressions. Our findings thus uncover species-specific oxygen sensing as a unifying mechanism for limb regeneration initiation across vertebrates, reveal how aquatic subatmospheric habitats may enhance regenerative capabilities, and identify targetable barriers to unlock latent limb regenerative programs in adult mammals.

## Introduction

Unlike terrestrial mammals, amphibians that thrive in aquatic subatmospheric oxygen environments can possess the remarkable ability to regenerate limbs (*1*, *2*). Following limb amputation, the large injured area comes into direct contact with the environment, and regeneration begins with basal epithelial cells covering the amputation plane and the formation of regenerative populations such as a specialized wound epidermis, and a blastema (*3*). The blastema contains muscle stem cells and multipotent connective tissue (CT) progenitors to repopulate limb cell types including fibroblast and cartilage (*4–8*). In contrast, mammals cannot rapidly heal amputation wounds (*9*). Numerous hypotheses exist aiming to explain the mammalian inability to regenerate limbs, such as failure to form a specialized wound epidermis or CT progenitors, the evolution of thermoregulation, and epigenetic alterations (*10–17*). However, these have not resulted in definitive evidence demonstrating a latent capacity for limb regeneration in mammals. Thus, whether, and how, mammals can activate limb regenerative processes after amputation, and how environmental conditions mechanistically shape regenerative capacities across species remain unresolved.

The apical ectodermal ridge (AER), a transient signalling center population controlling limb development (*18*), was recently identified as the single-cell identity constituting the specialized wound epidermis in regenerating *Xenopus laevis* tadpole limbs (*19*). While these aquatic tadpoles can rapidly heal wounds throughout their life (*1*, *20*, *21*), they lose their regenerative capacity towards metamorphosis and become unable to re-form AER cells after amputation (*19*, *21*, *22*). Given that mammals form AER cells during limb development (*18*), they may retain the molecular and cellular foundation to initiate limb regenerative processes following amputation.

Here, using embryonic mouse and *Xenopus* tadpole models combined with imaging, single-cell-omics, histone modification profiling, and cross-species comparisons, we unveil how environmental oxygen levels and species-specific oxygen-sensing orchestrate regenerative abilities across vertebrates. Our findings reveal that oxygen-sensing regulates the initiation of limb regeneration by directing cellular mechanical properties during wound healing, and establishing a regenerative histone landscape and metabolic state, linking environmental and genetic features to latent regenerative abilities. In doing so, we identify species-specific oxygen-sensing capacity as a fundamental mechanism underlying differences in regenerative abilities across vertebrates, offering insights into the evolution of regeneration and paving the way toward adult mammalian limb regeneration.

## Results

### Embryonic mouse limb explants can perform rapid wound healing depending on the culture conditions

Previously, we demonstrated that Nieuwkoop and Faber (NF) stage 53-54 (*23*) *Xenopus* tadpole limbs can initiate regeneration *ex vivo* in serum-containing media in a 96-well plate (*19*). In this setting, explants undergo rapid wound healing and re-form AER cells at distal amputation planes **(Fig 1A, Fig S1A),** allowing close examination of regeneration initiation in a highly controllable environment stripped from the bodily influences. To determine whether mammalian limbs possess a latent ability to initiate regeneration *ex vivo*, we examined E12.5 mouse limbs, which share similar morphology as NF stage 53-54 *Xenopus* limbs **(Fig S1B),** and show similar single-cell transcriptional programs (*24*). E12.5 limbs can mimic development *ex vivo* in serum-free media in an air-liquid interface (ALI) culture, where approximately half the sample is exposed to air (*25*) **(Fig 1B, Fig S1C)**, providing a suitable culture condition to study regeneration potential. Despite these features, we found that amputations to E12.5 mouse limb explants did not result in wound closure; epithelial cells remained at the sides or internalized, and had excessive chondrogenesis **(Fig 1C, Fig S2A)**. Performing limb amputations to *ex utero* grown E12.5 whole mouse embryos yielded similar results, suggesting that the failure to perform rapid wound healing is not due to disconnection from the body, although this cannot be ruled out entirely since embryos showed signs of suboptimal development *ex utero* **(Fig 1D, Fig S2B-C)**.

**Figure 1.**
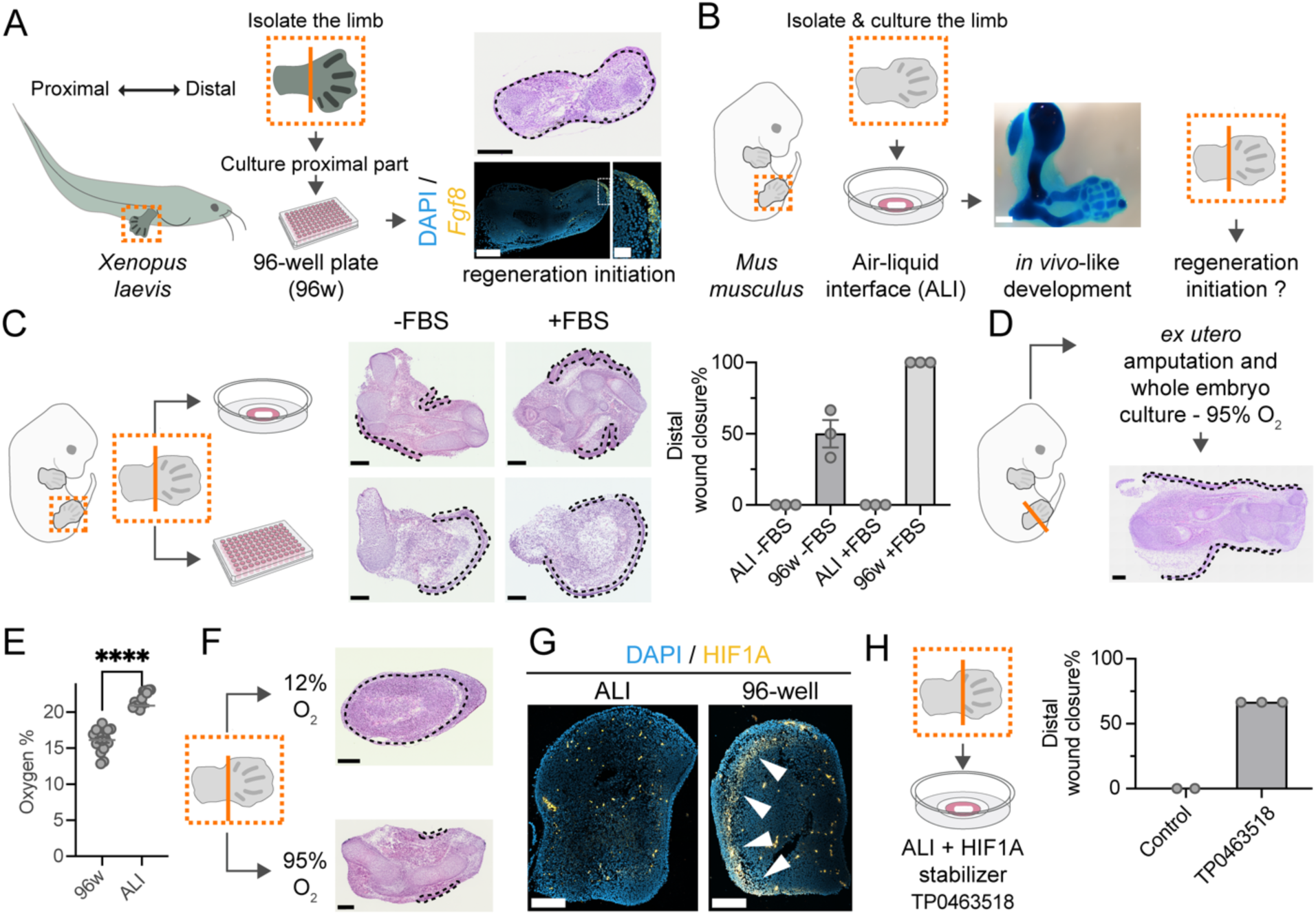
Subatmospheric oxygen promotes widespread wound healing in amputated embryonic mouse limbs, partly by HIF1A stabilization. **(A)** (Left) Schematic representation of the generation of regenerating *Xenopus* tadpole limb explants, where hindlimbs were isolated and amputated at the ankle depicted by an orange line. The proximal part of the amputated limb is cultured *ex vivo* in 96-well plate (96w). (Right-Top) Representative histology image of a regenerating *Xenopus* explant at 3 days post-amputation (dpa), with dashed lines outlining the basement membrane. Scale bar = 200 μm. (Right-Bottom) Representative confocal HCR image of *Fgf8* mRNA in *Xenopus* 3 dpa explant distal skin. Dashed rectangle highlights zoomed *Fgf8*+ skin cells. Blue: DAPI; Yellow: *Fgf8* mRNA. Scale bar = 250 μm; (zoomed image) 50 μm. **(B)** (Left) Schematic of mouse developing limb explant preparation, showing isolated intact limbs cultured in air-liquid-interface (ALI). (Right) Representative image of a whole-mount alcian blue staining of E12.5 mouse limbs after 5 days in ALI culture. Scale bar = 1000 μm. **(C)** (Left) Schematic of experimental design to test the regenerative potential of mouse limb explants cultured in 96w or ALI, with or without serum, FBS. (Middle) Representative histology images of mouse limb explants at 3 dpa under different culture conditions. Dashed lines surround epithelial cells. Scale bar = 200 μm. (Right) Bar plot quantifying the percentage of explants with distal wound closure at 3 dpa under the indicated conditions. Each dot represents an individual experiment. ALI-FBS: N=3, total n=0/10 explants closed distal wound area; 96w -FBS: N=3, total n=5/10 explants closed distal wound area; ALI +FBS: N=4, total n=0/11 explants closed distal wound area; 96w +FBS: N=4, total n=12/12 explants closed distal wound area. **(D)** (Left) Schematic illustrating *ex utero*-grown whole E12.5 mouse embryo cultures used to assess hindlimb postamputation response. (Right) Representative histological image of a 2 dpa limb, with dashed lines surrounding epithelial cells. N = 1, n = 0/3 embryos closed distal wound area. Scale bar = 200 μm. **(E)** Scatter plot of oxygen levels in the culture media of mouse limb explants in 96w or ALI over 3 days. Mouse ALI N = 3, total n = 8; Mouse 96w N = 3, total n = 17. **(F)** Representative images of mouse explants under controlled 12% and 95% oxygen levels. Dashed lines indicate the basement membrane (top image) and epithelial cells (bottom image). 12% oxygen N=3, total n=8/8 explants closed distal wound area; 95% oxygen N = 3, total n=0/9 explants closed distal wound area. Scale bar = 200 μm. **(G)** Representative confocal images of HIF1A immunofluorescence in mouse explant sections at 6 hours post-amputation (hpa) cultured in 96w or ALI. N=2, n=6 for both conditions. White arrows show HIF1A enriched areas. Scale bar = 250 μm. **(H)** Bar plot showing the percentage of mouse explants with distal wound closure at 3 dpa in ALI upon TP0463518 treatment. Vehicle treatment is used as the control. Each dot represents an individual experiment. Vehicle control N = 2, total n=0/9 explants closed distal wound area; TP0463518 N=3, total n=6/9 explants closed distal wound area.

To assess if culture conditions influence differences between mouse and *Xenopus*, we cultured mouse explants in serum-free media in 96-well plates **(Fig 1C)**. Unexpectedly, basal epithelial cells covered the distal amputation plane in half of the explants, with decreased underlying chondrogenic cells **(Fig 1C)**, and the inclusion of serum resulted in wound healing and chondrogenic suppression in all samples **(Fig 1C),** mimicking the *Xenopus* response. In contrast, even with serum, none of the ALI samples exhibited similar wound healing **(Fig 1C),** demonstrating that culture conditions critically impact the wound healing response.

### Subatmospheric oxygen and HIF1A stabilization induce rapid wound healing in embryonic mouse explants

Because oxygen diffusion rates vary with the culture surface area (*26*), and since parts of explants in ALI are directly exposed to atmospheric oxygen, we hypothesized that environmental oxygen influences wound healing. Using a sensitive fiber-optic oxygen sensor, we first determined that mouse and *Xenopus* explant media in 96- well plates, as well as our *Xenopus* tadpole husbandry tanks had 12–17% subatmospheric oxygen levels **(Fig 1E, S3)**. Meanwhile, ALI cultures had 21% oxygen **(Fig 1E)**, and *ex utero* cultures required 95% oxygen **(Methods),** suggesting subatmospheric oxygen could be promoting initiation of regeneration.

To directly test the effect of oxygen, we exposed mouse explants to 12% oxygen (**Methods**), which induced a remarkable wound healing response in all explants: they were covered with skin cells with strong chondrogenic suppression **(Fig 1F)**. In contrast, under the same setting, but with 95% oxygen, no wound healing occurred **(Fig 1F)**, although this level of oxygen was not impairing limb morphogenesis **(Fig S4A)**, confirming that environmental oxygen plays a key role in the execution of rapid wound healing.

Low oxygen can stabilize the oxygen-sensitive HIF1A, known to enhance wound healing following certain minor injuries (*27*, *28*). However, it was unclear if it can also act upon limb amputation, which poses a large and complex wound area. Examining this, we first found more HIF1A+ cells in explants in 96-well plate or 12% oxygen, and their notable absence in ALI or 95% oxygen samples (**Fig 1G, Fig S4B**). Treating explants in ALI with HIF1A stabilizers (DMOG, TP0463518 or IOX2) further supported the involvement of HIF1A, as TP0463518 overcame the negative effect of atmospheric oxygen, and had notable increase in number of HIF1A+ cells **(Fig 1H, Fig S4C-D).** Together, these results indicate that subatmospheric oxygen levels promote rapid wound healing following limb amputation partly through HIF1A stabilization.

### Subatmospheric oxygen changes cell shape and mechanical features during wound healing

We next explored how oxygen levels alter wound healing at the molecular and cellular levels. In intact limbs, *Xenopus* basal epithelial stem cells displayed apical actin enrichment (**Fig 2A, 2F**). Within six hours post amputation, however, migrating leading edge cells elongated, their actin filaments showed a more uniform distribution, and there was little actin accumulation towards the basement membrane (**Fig 2B, 2F**). Regeneration incompetent tadpoles, which also live in subatmospheric oxygen and exhibit rapid wound healing, showed a similar trend, except that their basal cells had more pronounced cell elongation and less apico-basal actin enrichment upon amputation (**Fig S5A-B**), indicating changes in cell mechanics and flexibility in cytoskeletal arrangements.

**Figure 2.**
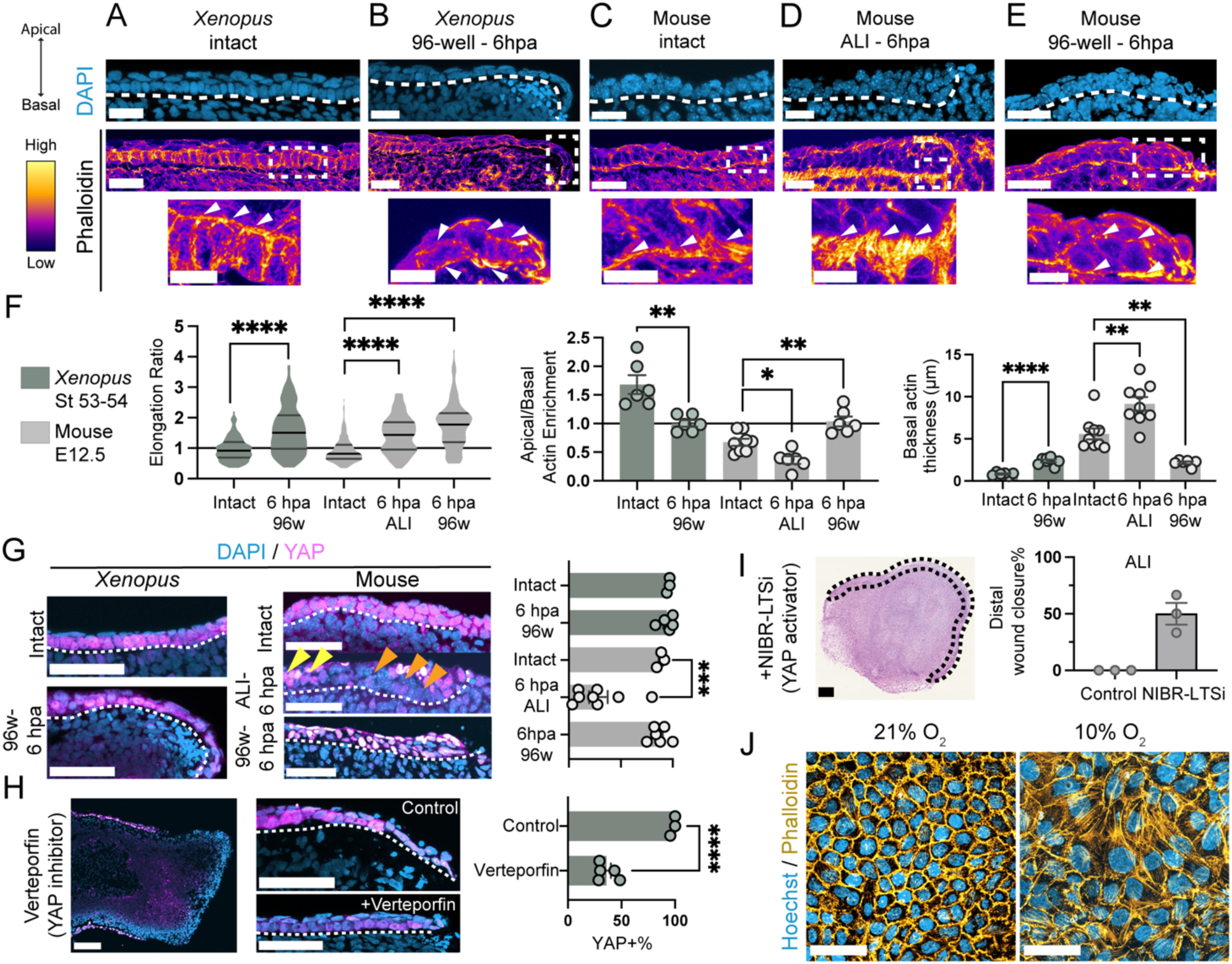
Subatmospheric oxygen culture conditions alter cell shape, actin dynamics, and YAP activity, enabling rapid wound healing. **(A-E)** (Top) Confocal images of sectioned limb basal epithelial cells of *Xenopus* and mouse intact or 6 hours post-amputation (hpa) under 96w or ALI culture. Dashed line labels basement membrane. Blue = DAPI. (Middle) Phalloidin (actin filaments)-stained confocal images of *Xenopus* and mouse limb sections under the same conditions. Heat map intensity colours via LUT in Fiji. (Bottom) Zoomed regions (dashed rectangle) highlighting actin levels. White arrows indicate regions of actin enrichment or depletion. Scale bar top and middle rows = 25 μm; zoomed images bottom row = 10 μm. N = 3 for all conditions, Xenopus intact total n = 6; Xenopus 96w 6 hpa total n = 7; Mouse intact total n= 8; Mouse ALI 6 hpa total n= 6; Mouse 96w 6 hpa total n= 6. **(F)** (Left) Violin plot quantifying the elongation ratio of individual basal epithelial cells in *Xenopus* and mouse limbs under indicated conditions. (Middle) Bar plot showing apical-to-basal actin enrichment ratios in group of basal epithelial cells. Each dot represents individual sections. (Right) Bar plot of basal actin thickness in basal epithelial cells. Each dot represents individual sections. Xenopus intact N=3, total n = 6, total quantified cell n = 119; Xenopus 96w 6hpa N = 3, total n= 7, total quantified cell n = 139; Mouse intact N= 3, total n= 8, total quantified cell n= 159; Mouse 6hpa ALI N= 3, total n=6, total quantified cell n= 120; Mouse 96w 6hpa N= 3, total n= 6, total quantified cell n= 119. Total quantified cell number applies to elongation ratio. **(G)** Representative active nuclear YAP immunofluorescence confocal images of basal epithelial cells in *Xenopus* and mouse explants under indicated conditions. Dashed lines outline the basement membrane. Yellow arrows mark nuclear nuclear YAP in proximal cells, and orange arrows highlight its absence in distal basal epithelial cells. Scale bar = 50 μm. (Right) Bar plot quantifying distal basal epithelial cells with nuclear YAP. Xenopus intact N = 3, n = 3; Xenopus 96w 6hpa N = 3, n=5; Mouse intact N= 3, n=3; Mouse ALI 6hpa N=3, n=8; Mouse 96w 6hpa N= 3, n=6. **(H)** (Left) Representative active nuclear YAP immunofluorescence confocal image of amputated *Xenopus* limbs treated with Verteporfin, showing lack of wound closure at 3 dpa and lack of epithelial cells at the distal limb. Vehicle control N= 3, total n= 9/9 explants closed distal wound area, Verteporfin N = 3, n=0/8 explants closed distal wound area. Scale bar = 100 μm. (Middle) Representative confocal images of control or Verteporfin-treated *Xenopus* 96w limbs at 6 hpa. Scale bar = 50 μm. (Right) Bar plot quantifying basal epithelial cells with nuclear YAP after Verteporfin treatment at 6 hpa. Vehicle control N= 3, n=9, Verteporfin N = 3, n=9. **(I)** (Left) Representative histology image of a mouse limb explant in ALI treated with the YAP activator NIBR- LTSi, showing distal wound closure at 3 dpa. Scale bar = 200 μm. (Right) Bar plot of explants with distal wound closure upon NIBR-LTSi treatment in ALI. Each dot represents an individual experiment. Vehicle N = 3, total n=0/9 explants closed distal wound area; NIBR-LTSi N=3, total n=4/9 explants closed distal wound area. **(J)** Representative confocal images of Phalloidin-stained MDCK cells cultured at 21% and 10% environmental oxygen. Blue = Hoechst; Yellow = Phalloidin. 21% oxygen N = 6, 10% oxygen N = 3, individual wells total n=8. Scale bar = 50 μm.

Unlike *Xenopus*, basal epithelial cells in the intact mouse limb showed not apical, but basal actin enrichment (**Figure 2C, 2F**). After amputation, basal cells in ALI showed a striking increase in actin accumulation towards the basement membrane, and distal potential migrating cells appeared to be blocked in movement by this accumulation and other populations (**Figure 2D, 2F**). Interestingly, however, mouse cells in 96-well plates or exposed to 12% oxygen had more uniformly distributed apico-basal actin and significantly reduced actin accumulation (**Fig 2E, Fig S6A-B)**, mirroring the *Xenopus* response. In both ALI and 96-well plate, mouse basal cells showed cell elongation following amputation (**Fig 2D-E)**. In contrast to these, 95% oxygen showed less cell elongation, more uniform apico-basal actin enrichment and moderate actin accumulation (**Fig. S6A-B**).

We also found distal leading edge basal cells in ALI or 95% oxygen samples to have increased LAMINA/C, which is enriched in stiffer nuclei (*29*), whereas 96-well plates or 12% oxygen levels had less enrichment (**Fig. S6C**). Thus, we concluded that environmental oxygen modifies cell mechanical properties, illustrated by changes in cell shape, actin organization, and nuclear rigidity of basal epithelial cells near the amputation plane, in a manner that aligns with it influencing rapid wound healing. Moreover, because these pronounced changes occur within six hours after amputation—even though the *Xenopus* and mouse intact limb cells display similar single-cell transcriptomes (*24*)—our results suggest that post-transcriptional mechanisms may mediate the rapid wound healing we observe.

### Subatmospheric oxygen promotes YAP activation, which is necessary and sufficient for rapid wound healing after amputation

Having seen cellular biomechanical changes due to elevated oxygen levels, we asked if oxygen alters post-transcriptionally regulated mechanosensing by YAP activation (*30*, *31*), which is critical for skin wound healing (*32*) and *Xenopus* limb regeneration (*33*).

Basal epithelial cells in both regeneration-competent (**Fig 2G**) and -incompetent *Xenopus* tadpoles (**Fig S5C)** had nuclear active YAP before and after amputation. However, surprisingly, mouse basal epithelial cells in ALI or 95% oxygen following amputation had significantly reduced nuclear YAP compared to intact limbs, particularly in the distal cells (**Fig 2G, S6D)**. Meanwhile, mouse basal cells in 96-well plates or under 12% oxygen retained nuclear YAP, suggesting subatmospheric environmental oxygen sustains YAP activation (**Fig 2G, S6D**). Together, these results uncovered that only a small number of distal basal epithelial cells close to the amputation plane show potential YAP-related biomechanical changes in mice, regulated by environmental oxygen levels.

To dissect the link between environmental oxygen and YAP activation, we treated *Xenopus* explants in 96-well plates with Verteporfin to impair YAP activity (*34*). Verteporfin treatment resulted in a complete lack of wound healing in all explants and a loss of distal but not proximal basal epithelial cells, presumably due to apoptosis (**Fig 2H**). In contrast, treating mouse explants in ALI with NIBR-LTSi, a LATS kinase inhibitor enabling YAP activation (*35*), induced wound healing in some explants (**Fig 2I, Fig S6E**), suggesting that active YAP can overcome the negative effect of atmospheric oxygen, and operates downstream of the effect of environmental oxygen.

Cell shape changes, YAP activation, and wound healing in mice and *Drosophila* (*36–38*) have been linked to solid-to-fluid-like tissue transition, a biomechanical property-driven mechanism for cellular mobility (*36*). Therefore, our results raise the possibility that oxygen modulates the solid-to-fluid-like tissue transition, complementing the established notion of hypoxia (<5% O_2_) promoting cell migration and wound healing (*39*, *40*). Using MDCK epithelial monolayers, a model for studying the solid-to-fluid-like transition (*41–43*), we observed that atmospheric oxygen (21% oxygen), produced a honeycomb, solid-like cell arrangement with actin-enriched borders (**Fig 2J, Fig S7**). Lowering oxygen to 10% induced cell elongation, increased neighbours per cell, enlarged nuclei (**Fig 2J, Fig S7**), and reduced actin enrichment at borders - consistent with a solid-to-fluid-like tissue transition behaviour (*44*), although this phenotype also aligns with properties of epithelial-mesenchymal-transition (*45*). Thus, we concluded that subatmospheric oxygen alters cellular properties into such indicative of fluid-like tissue, though live-imaging-based assessment of basal epithelial cells is warranted to determine whether subatmospheric oxygen can directly promote tissue fluidization.

### Subatmospheric oxygen allows induction of a regenerative apical-ectodermal-ridge (AER) and blastema gene expression in embryonic mouse limbs

Next, we asked if mouse explants could advance in full regeneration initiation and re-form AER cells, a prerequisite for blastema formation (*46*). Examining 3 dpa 96-well plate explants for epithelial *Fgf8* expression— a marker for AER—we could not detect AER formation **(Fig 3A)**. Thus, while subatmospheric oxygen enhances wound healing, it is insufficient for full initiation of regeneration (*16*, *47*).

**Figure 3.**
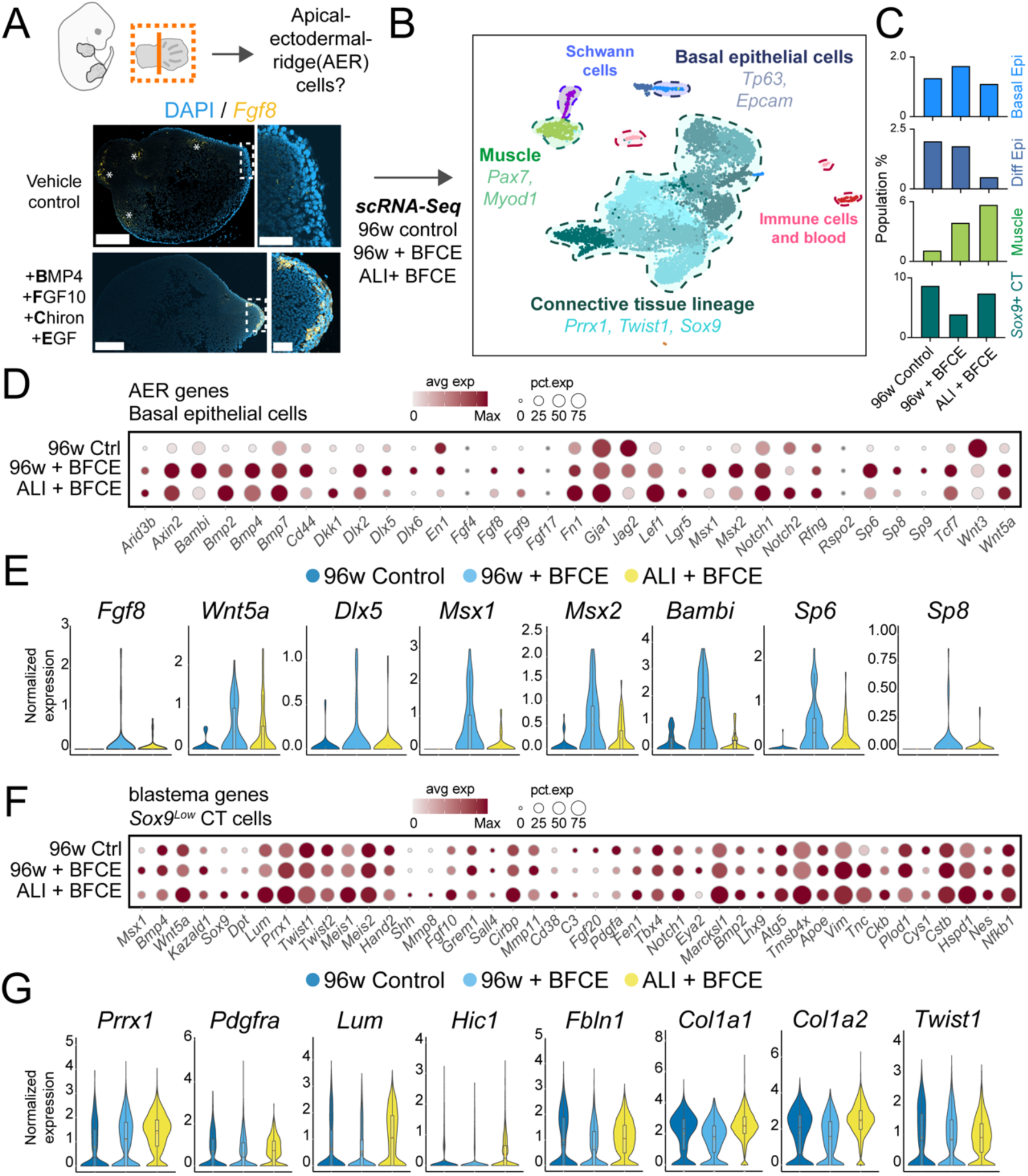
Amputated embryonic mouse limbs exhibit full initiation of regeneration upon exogenous supplementation in subatmospheric environments. **(A)** Representative confocal HCR images for *Fgf8 mRNA*, a marker for regenerative apical ectodermal ridge (AER) cells in (top) no treatment or vehicle control and (bottom) BMP4, FGF10, Chiron, EGF (BFCE) treated explants in 96w. Dashed squares indicate magnified images on the right. * indicates non-specific staining. N = 2, n = 6. Scale bar = 250 μm for full sections, and 50 μm for zoomed images. **(B)** Integrated UMAP representation of 3 dpa scRNA-Seq samples from (1) 96w with vehicle control, (2) 96w treated with BFCE, and (3) ALI treated with BFCE. Full cluster annotation is available in Fig S8. **(C)** Bar plots showing changes in the percentage of indicated cell clusters across experimental groups. *Sox9+* connective tissue (CT) cells represent cartilage. **(D)** Dot plot showing AER-associated gene expression in basal epithelial cells across the three experimental conditions. **(E)** Violin plots of representative expression levels of key AER genes in basal epithelial cells across the three experimental conditions. **(F)** Dot plot showing blastema-associated gene expression in *Sox9^Low^* (non-cartilage) CT lineage cells across the three conditions. **(G)** Violin plots of fibroblast-specific gene expression levels in *Sox9^Low^* (non-cartilage) CT lineage cells across the three conditions.

This outcome parallels regeneration-incompetent *Xenopus* tadpoles, which can heal rapidly but fail to re-form AER cells post-amputation (*19*). In these tadpoles, chondrocyte-derived inhibitors (*19*) and insufficient signals from connective tissue (CT) progenitors create a non-permissive niche for AER re-formation (*16*, *47*). Given that amputated E12.5 mouse limbs show robust chondrogenesis and more differentiated CT (**Fig 1C-D**), we hypothesized that their limb environment similarly lacks necessary ligands for AER induction.

To overcome this barrier, we treated explants with a cocktail of AER-inducing factors—BMP4, FGF10, the *Wnt* pathway activator CHIR99021 (Chiron), and EGF (hereafter, BFCE) (*47*, *48*) **(Fig 3A)**. Strikingly, BFCE treatment induced the AER marker *Fgf8* in basal epithelial cells in 96-well plates (**Fig 3A**). To comprehensively characterize this response, we performed scRNA-Seq (**Fig 3B-C, Fig S8)** which remarkably revealed that key AER genes were induced in basal epithelial cells (e.g., *Fgf8/9, Bmp2/4/7*, *Wnt5a*, *Notch1*, *Tgfb1*, *Dlx2/5/6*, *Msx1/2, Sp6/8, Cd44*) (**Fig 3D-E)**, despite their low abundance in limbs and commonly observed problems with capturing AER cells using scRNA-Seq (*49–52*). In contrast, the vehicle-control lacked AER signatures (**Fig 3C-E)**, and had more *Sox9^High^* CT (chondrogenic) populations **(Fig 3C, Fig S8E, Supplemental Table 1)**, supporting the *Xenopus*-based idea that cartilage-derived factors suppress AER re-formation (*19*).

We explored CT and muscle lineage that would interact with re-formed AER cells. E12.5 limbs already express significant limb developmental programs, which may mask the impact of AER cells. Nonetheless, CT cells in 96-well plate BFCE samples had significantly increased expression levels of limb development and blastema genes (e.g., *Msx1, Grem1, Prrx1*, *Kazald1, Lhx9, Mmp11, Nes, Tnc, Plod1* (*24*, *53*)) (**Fig 3F, Fig S9A)**, and displayed fewer chondrogenic cells (**Fig 3C, Supplemental Table 1)** and fibroblastic markers (**Fig 3G-H)**. These features suggest a strong shift towards multipotent blastema CT progenitors. Additionally, BFCE treated explants had more muscle cells (**Fig 3C, Supplemental Table 1**). Thus, latent regenerative programs and cell fates can be indeed activated under subatmospheric oxygen in amputated embryonic mouse limbs.

To test whether these signatures can be detected under atmospheric oxygen, we performed scRNA-seq on BFCE-treated mouse explants in ALI (**Fig 3B, Fig S8**). Compared to the low-oxygen condition, ALI samples contained fewer basal epithelial cells (**Fig 3C, Supplemental Table 1)** with significantly weaker AER gene expression (e.g., *Msx1, Sp6, Bambi*) (**Fig 3E**) and CT-associated *Fgf10* and abundant *Shh* expression (**Fig S9B**). Although CT cells still showed substantial blastema gene expression (**Fig 3F, Fig S9A**), they had higher cartilage (**Fig 3C, Supplemental Table 1**), and fibroblastic signatures (**Fig. 3G**). Thus, BFCE treatment in ALI led to fewer cells with multipotent CT progenitor properties. In sum, atmospheric oxygen is not compatible with the BFCE treatment for establishing a regenerative state in a mammalian context.

Lastly, unlike *Xenopus*, another limb regeneration-competent amphibian, axolotls induce key AER genes in blastema CT cells (*24*, *54*, *55*). We found no similar axolotl-like AER gene expression in CT cells under any mouse condition (**Fig. S9C**), suggesting that mouse regenerative signatures more closely parallel those of *Xenopus*, in line with their commonalities for their limb development (*56*).

### Atmospheric oxygen does not substantially alter chromatin accessibility for AER gene regions

Our results indicated that subatmospheric oxygen enhances AER gene expression. We thus hypothesized that atmospheric oxygen might impair the chromatin accessibility necessary for AER induction. To test this, we performed time-course single-cell Multiome (scMultiome, combined single-nucleus ATAC-seq and RNA-seq) on intact mouse embryonic limbs at baseline and at 6 hours, 1 day, and 3 days post-amputation **(Fig 4A, Fig S10)**. As expected, intact E12.5 limbs expressed AER genes, likely in forming digit tip skin (*18*), which disappeared after amputation **(Fig S10F)**. Moreover, consistent with our histology data **(Fig 1C)**, the scMultiome analysis detected a sharp increase in *Sox9^High^* CT chondrogenic cell numbers **(Fig S10F)**, validating our dataset.

**Figure 4.**
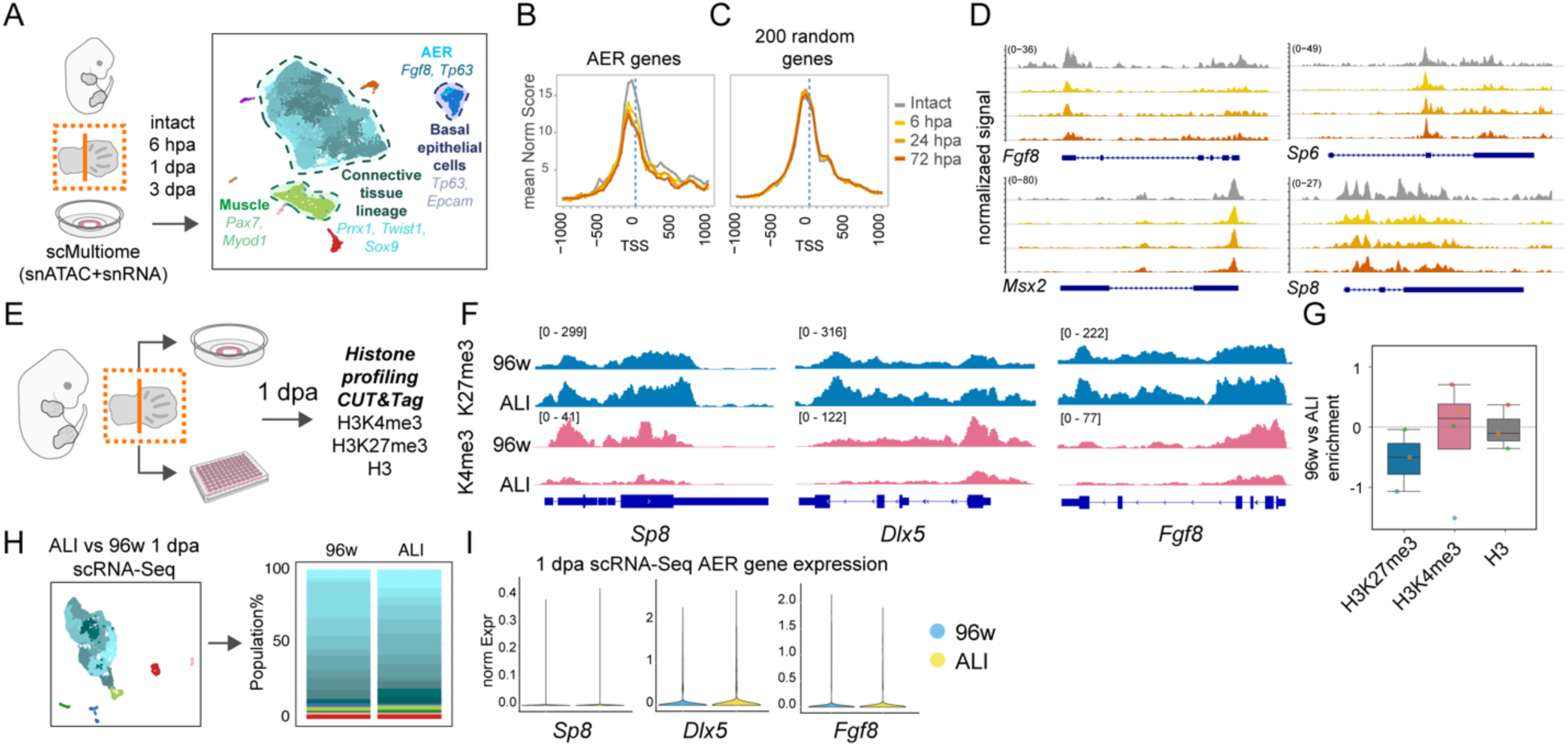
Subatmospheric oxygen promotes a histone landscape favoring regenerative gene expression in embryonic mouse limbs. **(A)** RNA only UMAP representation of scMultiomics (single-nucleus ATAC-seq (snATAC-Seq) + single-nucleus RNA-seq (snRNA-Seq)) data from embryonic mouse limbs cultured in ALI, including intact E12.5 limbs, 6 hpa, 1 dpa, and 3 dpa samples. Full cluster annotation is available in Fig S10. **(B)** Enrichment plot of snATAC-Seq data for combined basal epithelial and AER clusters, showing chromatin accessibility of key AER genes. Plotted time points are colour coded and shown on the right. **(C)** Enrichment plot of snATAC-Seq data for combined basal epithelial and AER clusters, showing chromatin accessibility of 200 random genes as a control. Plotted time points are colour coded and shown in Fig 4B. **(D)** Example time-course snATAC-Seq genome browser snapshots showing accessibility dynamics for AER genes in combined basal epithelial and AER cell clusters. **(E)** Schematic describing the experimental setup for CUT&Tag histone profiling of mouse explants. **(F)** Genome browser snapshots of representative replicates showing the enrichment of histone marks (H3K4me3—enriched at active genes; H3K27me3—enriched at repressed genes) at key AER genes. **(G)** Box plot comparing the ratio of enrichment for H3K4me3, H3K27me3, and total H3 modifications between 96w and ALI conditions for all TSS peaks. **(H)** UMAP representation of integrated scRNA-seq data from mouse embryonic limbs at 1 dpa in ALI and 96w. (Right) Bar plot showing cell population percentages derived from scRNA-seq data. Full cluster annotation is available in Fig S14. **(I)** Violin plots showing low expression levels of selected AER genes in 1 dpa limbs cultured in 96w and ALI.

Contrary to our initial hypothesis, atmospheric oxygen did not reduce chromatin accessibility upon amputation; instead, most cell types showed gains of accessibility **(Fig S11A-B)**. Within AER and basal epithelial cell clusters, accessibility at key AER gene regions decreased only marginally over time **(Fig 4B-D, Fig S10C-D)**. Thus, atmospheric oxygen has a minor impact on chromatin accessibility, including at AER-related loci.

### Subatmospheric oxygen promotes a regenerative histone landscape by increasing H3K4me3 and decreasing H3K27me3 levels

Since atmospheric oxygen did not substantially restrict chromatin accessibility, we asked whether histone modifications might serve as an alternative mechanism priming regenerative gene expression under low oxygen. We cultured mouse explants in ALI or 96-well plate for 1 day and performed CUT&Tag against histone modifications H3K4me3 (active), and H3K27me3 (repressive), with H3 and spike-ins **(Fig 4E, Fig S12)**.

Strikingly, in most 96-well plate samples, critical AER-related gene regions were enriched for H3K4me3 and reduced for H3K27me3 compared to ALI **(Fig 4F, S12D)**, representing a priming mechanism for the superior effectiveness of BFCE treatment under subatmospheric oxygen. This response was exemplified by key AER transcription factors such as *Dlx5* and *Sp8*, and to a lesser extent by AER ligands including *Fgf8* and *Fgf9* **(Fig 4F, S13)**. Similarly, other regeneration-related loci, including positional information-related *HoxA* and *HoxD* clusters, showed comparable changes (**Fig. S12E-G**). Moreover, we detected a mild widespread increase of H3K4me3 in 96-well plate, and for H3K27me3 in ALI, while H3 was similarly distributed between samples **(Fig 4G)**.

To determine whether these histone modification changes correspond to the formation of specific cell fates, we performed scRNA-seq at 1 dpa for both ALI or 96-well plate samples (**Fig 4H, Fig S14**). Both conditions revealed highly similar cell type distributions, although ALI had more *Sox9^High^* chondrogenic CT cells **(Fig 4H, S14D, Supplemental Table 1)**. Despite this, even chondrogenic genes such as *Sox9*, and *Col2a1* showed similar enrichment for H3K4me3 and reduction for H3K27me3 **(Fig S15)**. Because the majority of AER and blastema genes are not associated with chondrocytes but show similar histone modification changes (**Fig 4F-G, Fig S12D-E, S13**), these results suggest that the widespread changes we observed occur in cell types beyond chondrocytes, though we cannot fully rule out cell-type-specific effects.

Notably, at 1 dpa, neither 96-well plate nor ALI samples robustly expressed AER and regeneration-associated genes (**Fig S16**), although they showed histone modification changes indicative of gene activation (increase in H3K4me3 and decrease in H3K27me3) (**Fig S12D-G**). Thus, our results suggest that subatmospheric oxygen promotes a regenerative histone landscape via priming the chromatin for regenerative gene activation.

### Xenopus tadpole limbs are less sensitive to elevated environmental oxygen levels during regeneration initiation

Our findings uncovered that subatmospheric oxygen can promote, and atmospheric oxygen can impair wound healing and regenerative response in mouse limbs. Nonetheless, urodele amphibians such as newts and the axolotl—which have distinct limb developmental programs compared to *Xenopus* and mice (*56*)—can still regenerate their limbs on land and even at high oxygen levels (*57*). We therefore asked whether elevated oxygen levels could counteract regenerative processes in *Xenopus*.

Critically, both *in vivo* and *ex vivo*, *Xenopus* limbs were still able to perform rapid wound healing above 60% oxygen (**Fig. 5A-B)**, underscoring that even with the involvement of bodily factors such as the immune system or neurons, tadpoles are less sensitive to environmental oxygen during regeneration initiation. Moreover, even at 65% oxygen levels, *Xenopus* basal epithelial cells continued to show cell shape changes, actin dynamics, and YAP activity similar to explants cultured in subatmospheric oxygen **(Fig 5C-E)**. Furthermore, performing CUT&Tag against H3K4me3, H3K27me3, and H3 in 1 dpa *Xenopus* explants in ALI or 96-well plate revealed no obvious widespread or targeted changes in AER and regenerative gene regions between the conditions **(Fig 5F-H, Fig S17, S18)**. In an alignment, in either ALI or at 65% oxygen, *Xenopus* explants re-formed potential epithelial *Fgf8*+ AER cells **(Fig 5I)**. Lastly, performing scRNA-Seq on 1 dpa *Xenopus* limbs in ALI and 96-well plate revealed highly similar cell type distribution and cycling cell profiles **(Fig 5J, Fig S19, S20, Supplemental Table 1)**. Therefore, our results, combined with previous findings showcasing *Xenopus* can develop with minimal oxygen (*58*), highlight their inherently low oxygen-sensing capacity across both high and low oxygen spectrums.

**Figure 5.**
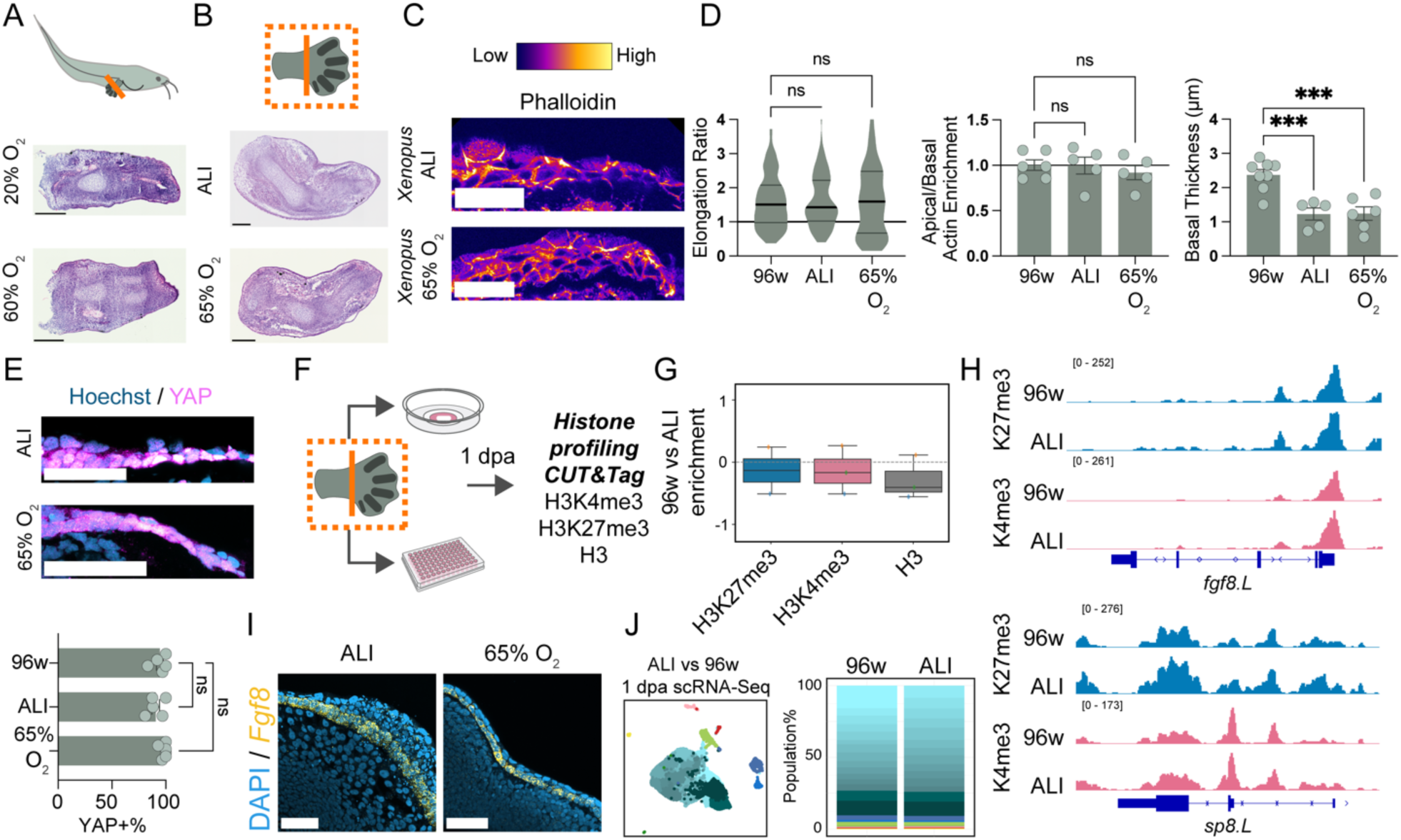
*Xenopus* limbs maintain actin dynamics, YAP activation, histone profiles, and cell type distribution despite elevated oxygen levels. **(A)** Representative 1 dpa histology image of an *in vivo* amputated *Xenopus* limb under 20% or 60% oxygen conditions. 20% oxygen N = 1, total n=4/4 tadpoles closed distal wound area; 60% oxygen, total n=8/8 tadpoles closed distal wound area. Scale bar = 200 μm. **(B)** Representative 3 dpa histology image of an *ex vivo* amputated *Xenopus* limb in ALI or under 65% oxygen conditions. ALI N = 3, total n=8/8 explants closed distal wound area; 65% oxygen N=3, total n=9/9 explants closed distal wound area. Scale bar = 200 μm. **(C)** Confocal images of phalloidin staining in *ex vivo* amputated *Xenopus* limbs at 6 hpa under ALI and controlled 65% oxygen conditions. Scale bar = 200 μm. **(D)** (Left) Violin plot quantifying the elongation ratio of individual basal epithelial cells in *Xenopus* limbs under indicated conditions. (Middle) Bar plot of apical-to-basal actin enrichment ratios for basal epithelial cells. Each dot represents individual sections. (Right) Bar plot of basal actin thickness in basal epithelial cells. Each dot represents individual sections. 96w N =3, total n = 7, quantified total cell n = 139; ALI N = 3, total n=5, quantified total cell n = 100; 65% oxygen N=3, total n=6, quantified total cell n=118. Total quantified cell number applies to elongation ratio. **(E)** Confocal images of active nuclear YAP staining in *ex vivo Xenopus* limbs at 6 hpa under ALI and controlled 65% oxygen conditions. (Bottom) Bar plots quantifying nuclear YAP in basal epithelial cells. 96w N = 3, n= 5, ALI N =3, n= 5; 65% oxygen N=3, n = 6. Scale bar = 50 μm. **(F)** Schematic representation of the experimental setup for CUT&Tag histone profiling in *Xenopus* explants. **(G)** Box plot comparing the ratio of enrichment for H3K4me3, H3K27me3, and total H3 modifications between 96w and ALI cultures for *Xenopus* limbs for all TSS peaks. **(H)** Genome browser snapshots of representative replicates for AER marker *Fgf8* and *Sp8* for H3K4me3, H3K27me3. **(I)** Confocal HCR image for *Fgf8* mRNA in 3 dpa amputated *Xenopus* limb distal skin under ALI and controlled 65% oxygen conditions. N = 3, n = 9. Scale bar = 50 μm. **(J)** UMAP representation of integrated scRNA-seq data from *Xenopus* limbs at 1 dpa under ALI and 96w. (Right) Bar plot showing cell population percentages derived from scRNA-seq data. Full cluster annotation is available in Fig S19.

### Species-specific oxygen-sensing capacity through regulation of HIF1A stability is a critical factor associated with limb regenerative competencies

Because our results strongly demonstrated that mouse and *Xenopus* exhibit different oxygen-sensing capacities which can alter regenerative potential, we sought to uncover the mechanism behind this difference, and to examine if this feature could explain regenerative competencies across species, including humans.

Oxygen-sensor effector transcription factor HIF1A is affected by oxygen levels through PHD enzymes (PHD1/2/3 encoded by *Egln2/Egln1/Egln3*) that hydroxylate HIF1A, and the E3 ubiquitin ligase VHL. VHL then binds to hydroxylated HIF1A, leading to its degradation (**Fig 6A**); additionally, HIF1AN can block HIF1A transcriptional activity (*59*, *60*). Therefore, changes in the expression of HIF1A regulators, such as decreases in *Egln2* or *Vhl* would lower oxygen-sensing capacity and retain stable HIF1A. Thus, to assess species-specific oxygen-sensing capacities, we investigated whether expression of HIF1A regulators differs between regenerative amphibians and non-regenerative mammals.

**Figure 6.**
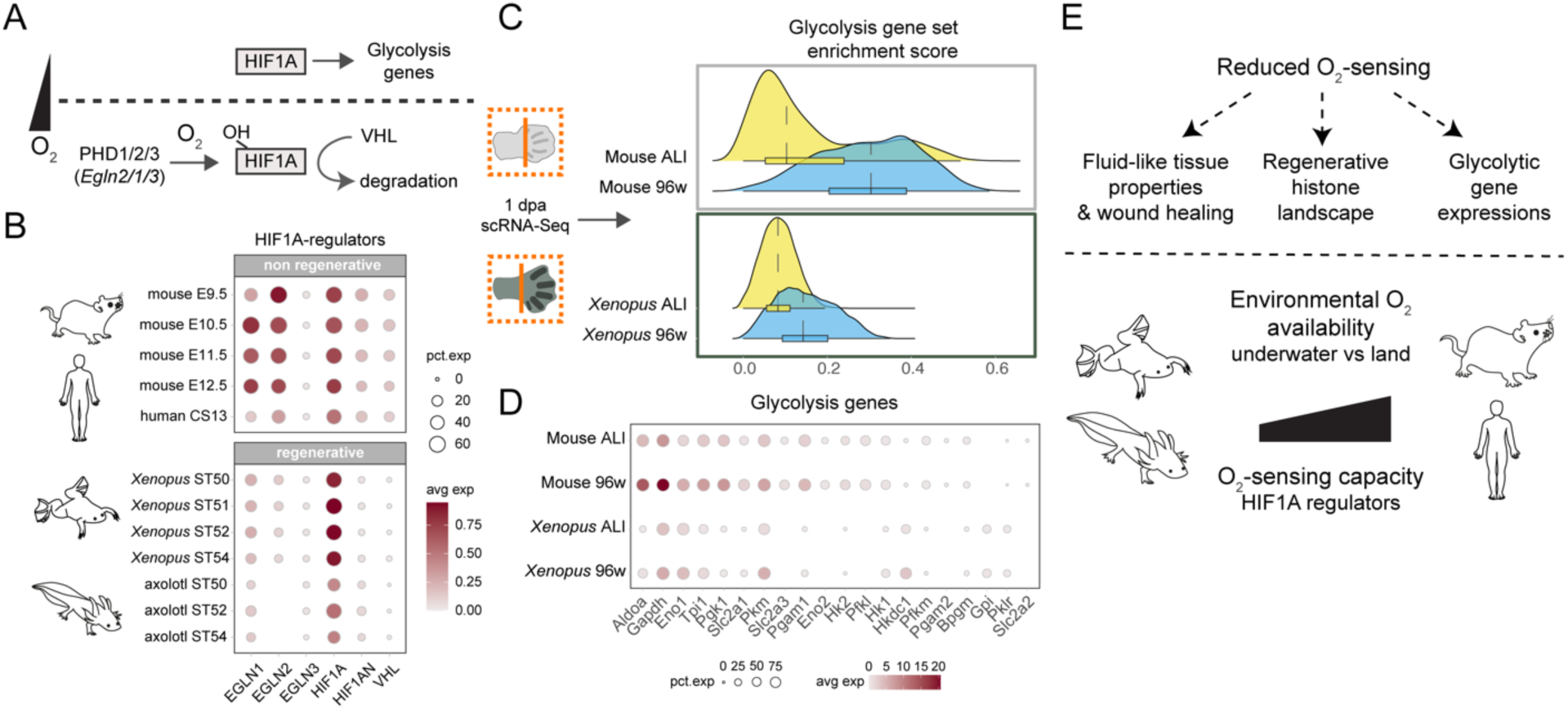
Regenerative amphibians, unlike non-regenerative mammals, exhibit reduced oxygen-sensing capacity and stable HIF1A activity illustrated by robust glycolytic gene expression. **(A)** Schematic representation of HIF1A regulation across varying environmental oxygen levels. **(B)** Dot plot showing gene expression profile of HIF1A and its regulators across publicly available *Xenopus*, axolotl, mouse, and human limb development scRNA-seq datasets. **(C)** Ridgeplot showing glycolysis gene set expression in 1 dpa scRNA-seq datasets for *Xenopus* and mouse limbs cultured in ALI or 96w. **(D)** Dot plot showing the expression levels of key glycolysis-related genes across regenerative and non-regenerative species. **(E)** Model illustrating critical regenerative processes influenced by environmental oxygen and species-specific oxygen-sensing capacities, and oxygen-sensing capacity diverging regenerative amphibians from non-regenerative mammals.

Excitingly, when we analyzed previously published *Xenopus* and axolotl limb development scRNA-seq datasets, we found them to have strikingly low expression levels of *Egln1*, *Egln2*, *Hif1an*, and *Vhl* compared to non-regenerative mouse and human limb development datasets **(Fig 6B)**. Moreover, *Xenopus* and axolotl limb regeneration datasets (*19*, *61*) showed similar phenotypes **(Fig S21A),** indicating conserved profiles beyond development.

Thus, regenerative amphibians show low oxygen-sensing capacities due to decreased expression of HIF1A regulators, potentially maintaining more stable HIF1A despite environmental changes. In contrast, non-regenerative mammals show higher HIF1A regulator expression and less stable HIF1A. Their oxygen-sensing capacity is thus enhanced, resulting in more pronounced responses to changes in oxygen levels.

### Regardless of environmental oxygen changes, Xenopus tadpole limbs robustly express glycolytic genes, a sign of stable HIF1A

Based on our results, we hypothesized that changes in environmental oxygen would elicit fewer molecular changes in *Xenopus* compared to mice. To test this, we interrogated gene expression signatures of varying oxygen-sensing capacities and focused on glycolysis genes, which are well-established to be associated with stabilized HIF1A (*59*, *62*). To compare the effect of oxygen and potential HIF1A stability in mouse and *Xenopus* in a highly controlled manner, we re-analyzed our scRNA-seq data from explants of both species at 1 dpa in 96- well plates (subatmospheric oxygen) and ALI (atmospheric oxygen) (**Fig 6C, S14, S19**).

Remarkably, *Xenopus* cells showed low but robust expression of key glycolytic genes irrespective of oxygen levels **(Fig 6C-D, S21B)**, validating their low oxygen-sensing capacity and stable HIF1A. In contrast, mouse cells in ALI compared to 96-well plate showed pronounced decreases in the expression of glycolysis genes (**Fig 6C-D, S21B)**, indicative of the less stable HIF1A, in agreement with our HIF1A staining results (**Fig 1G**). Both species exhibited cell-types-specific variation in glycolysis gene expression (**Fig S21C-D**), and enrichment in various metabolic gene sets, including oxidative phosphorylation, which had a small but significant increase under the ALI condition (**Fig S21C-E, Supplemental Table 2 and 3**). In conclusion, our highly controlled cross-species experimental platform further solidified the notion that species-specific oxygen-sensing capacities distinguish limb regenerative amphibians from non-regenerative mammals and predisposes them for successful initiation of regeneration.

## Discussion

Here we report that oxygen-sensing mechanisms are finely tuned across species, regulated by environmental oxygen and through HIFA regulator expression levels (**Fig 6E**). These differences, along with direct exposure to environmental oxygen following amputation, can either trigger or inhibit regenerative programmes by influencing cellular mechanics, histone modifications, glycolytic gene expression and rapid wound healing. This framework helps explain why *Xenopus* and numerous other animals from different clades inhabiting low oxygen aquatic environments display enhanced wound healing (e.g., zebrafish (*63*), dolphin (*64*)). Critically, low oxygen alone can promote wound healing in embryonic mouse limbs, and when coupled with AER-inducing signals, it can fully initiate limb regeneration in a mammalian context. Consequently, cross-species differences in oxygen sensing capacity emerge as a unifying principle demarcating regenerative from non-regenerative species.

Achieving mammalian limb regeneration may therefore demand holistic environmental interventions rather than targeting single lineages or signaling pathways. This aligns with studies that grafting limb bud mesoderm alone fails to induce limb regeneration in adult frogs (*65*) or indeed mice (*9*). While earlier research linked oxygen primarily to metabolic and proliferative changes (*66*, *67*), we now demonstrate its profound influence on cellular mechanics, and epigenetic states for limb regeneration. Although we focused on embryonic limbs, adults may face additional regeneration barriers, including oxygen-sensitive inflammatory or bone repair programs (*68–71*), both of which proposed to impede limb regeneration (*19*, *72*). Nevertheless, previous studies in adult mouse digit (*73*) and adult frog limb (*74*) have shown that short-term aquatic encapsulation of the freshly amputated area can lead to better regenerative outcomes, although these studies did not consider oxygen levels as a potential mechanism. Our results raise the possibility that these improvements derive, at least in part, from lowered local oxygen levels creating conditions more conducive to regeneration.

Subtle oxygen differences (96-well plate, 12-17% vs ALI, 21%) can drastically reshape cellular mechanics, and histone modifications, linked with morphogenesis and cell fate decisions, supporting early *physiological gradient theories* (*75*). Local oxygen gradients, in concert with species-specific and spatial HIF1A regulation, may influence not only regeneration but also disease, development, and evolutionary trajectories. These variations offer new insights into species comparisons, such as developmental tempo (*76–79*), and may enable more concrete analyses between mammals and distant species.

Our results in tadpoles, which exhibit low oxygen-sensing capacity and stable glycolytic gene expressions, suggest that high regenerative capacity might be retained at the expense of energy-intensive abilities such as thermoregulation, or enhanced brain function. Amphibians are also notably cancer-resistant (*80*, *81*). Counterintuitively, however, our identified molecular differences in oxygen-sensing in amphibians—lack of *Vhl* expression (*82*), persistent HIF1A stability (*83*), sustained glycolysis*—*are all known to promote cancer in mammals. This reinforces the idea that selective pressures for cancer resistance may be a more ancestral or separately evolved ability, allowing species to leverage regenerative capacities without uncontrolled cell growth.

Collectively, we have discovered that species-specific oxygen-sensing capacity is a fundamental, targetable principle that governs limb regeneration potential. Understanding cross-species oxygen-sensing differences brings us closer to unlocking latent limb regenerative capabilities in adult mammals and reframes our view of how environmental and evolutionary pressures sculpt regenerative diversity across species.

## Acknowledgements

We thank Life Science Editors for editing services; the EPFL Gene Expression Core Facility for their excellent support on 10X-Genomics and sequencing library preparations; the EPFL BioImaging and Optics Facility and the EPFL Histology Core facility for their technical assistance; the EPFL Center of PhenoGenomics for animal housing and mouse uterus dissections; and the European *Xenopus* Resource Centre (EXRC) for their services in supplying *Xenopus* embryos. C.A. is supported by EPFL School of Life Sciences ELISIR Scholarship, the Foundation Gabriella Giorgi-Cavaglieri, Branco Weiss Fellowship, SNSF NRP79 (407940–206349), and Novartis Foundation for Medical-Biological Research.

## Author contributions

Conceptualization: mainly C.A., G.T.; Experimentation: mainly G.T., K.H. performed CUT&Tag, A.E.D. helped with staining and performed image quantifications, C.A., F.Z., E.S., assisted experiments, A.V., and A.H. helped with whole mouse embryo culture and controlled oxygen experiments, L.N. and S.S designed and built the ex-utero embryo culture machine with the help of A.H., A.M. performed MDCK experiments, M.T. H.O. and T.K. performed *in vivo Xenopus* experiments; Computational analysis: M.L. performed all computational analyses except CUT&Tag, H.H. performed CUT&Tag analyses; Data interpretation: mainly C.A., G.T.; Funding acquisition: C.A.; Project administration: C.A.; Supervision: mainly C.A., F.Z. supervised CUT&Tag experiments; Writing – initial draft: mainly C.A., G.T.; Writing – review and editing: all authors

## Data and materials availability

All sequencing data and analysis scripts will be made available to reviewers on request and will be published upon publication. Requests for materials should be addressed to C.A.

## Materials and methods

### Animal experiments

All *Xenopus* and mouse explant experiments were performed at EPFL, Switzerland. Animal husbandry and experimental procedures adhered to the Swiss Federal Veterinary Office guidelines and were authorized by the Cantonal Veterinary Office (cantonal animal license no. VD3652c) and the national animal license (no. 33237). For these experiments: pregnant CD-1 mice were purchased from Charles River Laboratories, and outbred *Xenopus laevis* embryos were obtained from the European *Xenopus* Resource Centre (EXRC). All animals were housed in accredited husbandry facilities at EPFL, maintained at 22°C on a 12-hour light/dark cycle, and provided with species-specific appropriate diets. *Xenopus* tadpoles were group-housed in 13 x 26 mm tanks containing 3.6 L of standard Xenopus husbandry water. The quality of *Xenopus* water was regularly monitored and changed once a week to ensure optimal conditions.

Animal breeding, care, and experiments for *Xenopus in vivo* limb regeneration assays were performed in Yamagata University, Japan, under the approved protocols and guidelines of Animal Research Committee of Yamagata University (Approval Numbers: R4015 and R5062). For animal breeding, adult *Xenopus laevis* were obtained from Watanabe Breeding, and the embryos were generated by *in vitro* fertilization using standard protocols in 0.1X Marc’s Modified Ringer’s solution and maintained at 18°C.

### *Ex vivo* embryonic mouse limb development and post-amputation cultures in 96-well plates (96w) or air-liquid-interface (ALI)

The uteri from pregnant CD-1 mice were dissected and E12.5 embryos were isolated using fine forceps and scissors. Isolated embryos were transferred into 10 cm plates containing cold 1x PBS containing 1% Antibiotic-Antimycotic (ThermoFisher Scientific, 15240062).

For *ex vivo* embryonic mouse limb development experiments in Fig 1B and Fig S1C, the E12.5 mouse embryo hindlimbs were isolated from the animal body and placed in an air-liquid-interface (ALI). The ALI culture system was set as described previously (*25*), using Falcon organ culture dishes (ThermoFisher Scientific, 10049210). Briefly, 2 mL of BGJb media ((Thermo Fisher Scientific, 12591038), 1% Antibiotic-Antimycotic (Thermo Fisher Scientific, 15240062), 0.8 mM L-Glutamine (Thermo Fisher Scientific, 25030024), and 20% Fetal Bovine Serum Superior (FBS) (Sigma, S0615) (except experiments in Fig 1B-C, S1C, S4C, which did not contain FBS as indicated)) was placed in the center well of a 60 x 15 mm Falcon organ culture dish (ThermoFisher Scientific, 10049210). The limb was placed on top of a 1.0 µm PET Membrane (Corning, 353102) held by a stainless-steel grid section (ThermoFisher Scientific, 15330912) with an opening in the middle. In the outside well of the Falcon organ culture dish, 2 mL of 1x PBS containing 1% Antibiotic-Antimycotic was added to limit evaporation. These mouse limb explants were cultured in a cell culture incubator (Thermo Fisher Scientific, Heracell VIOS 160i SS) at 37°C with 5% CO₂. The media was changed every two days.

For *ex vivo* embryonic mouse limb post-amputation experiments, the isolated E12.5 mouse embryo hindlimb was additionally amputated (distal amputation) at the presumptive ankle, and the autopod was discarded. The remaining proximal part was placed on ALI with 2 mL or in a 96-well plate (Thermo Fisher Scientific, 167008) with 200 µL BGJb media (Thermo Fisher Scientific, 12591038) to examine distal post-amputation/regenerative response. No distinction was made between left and right hindlimbs. No sex determination was done for embryos.

### *Ex vivo Xenopus* tadpole limb post-amputation cultures in 96w or ALI

Tadpoles classified as regeneration-competent were at Nieuwkoop and Faber (NF) (*23*) stages ∼53–54, while regeneration-incompetent tadpoles were at NF stages ∼58–60, according to (*22*). *Xenopus* limb explant generation was performed as previously described (*84*). Briefly, tadpoles were anesthetized by submersion in 0.02% MS222 (Sigma Aldrich, A5040) in *Xenopus* husbandry water, and placed on a wet paper towel under a stereomicroscope. The hindlimb was isolated from the body. Hindlimb was amputated at the presumptive ankle (distal amputation) using fine scissors, and the autopod was discarded. The remaining proximal part was then transferred either to a 96-well plate with 200 µL of media or an air-liquid interface (ALI) chamber with 2 mL of L-15 media (L-15 (ThermoFisher Scientific, 21083027), 1% Antibiotic-Antimycotic, 0.8 mM L-Glutamine, and 20% FBS). No distinction was made between left and right hindlimbs. Sex determination was not possible for these tadpole stages. *Xenopus* limb explants were cultured in a temperature-controlled incubator (Memmert, HPP110eco) at 22°C without CO₂. The media was changed every two days.

### *Ex utero* whole mouse embryo cultures

For the *ex utero* whole mouse embryo culture and limb amputation experiment (Fig 1D, S2B-C), 12-day gestation CD1 pregnant female mice were euthanized using CO2, and embryos were dissected from the uterus. Then, the embryos were carefully dissected out of the yolk sac, with the placenta attached, while maintaining the integrity of the extra-embryonic membranes (amnion and yolk sac with the ectoplacental cone), as previously described (*85*, *86*) (Fig S2B). Once the embryo was exposed, the distal part of the right hindlimb (ankle level) was amputated at the presumptive ankle using scissors (Fig 1D). The embryos were then carefully transferred to glass bottles (Schmizo, 9000.42) containing 2 mL of BGJb media. The amputated embryos were allowed to develop in the rolling *ex utero* culture system for one or two days (as indicated in-text) at 37°C with 5% CO₂ and 95% O2, as indicated in Fig 1D, Fig S2.

### Rolling *ex utero* culture system

An in-house-built rolling *ex utero* culture system based on Hanna Lab *ex utero* culture system was used (*87*). This system consists of a gas mixer system (Oxystreamer, SH2CO) connected to a custom-built rotating bottle culture unit placed inside a 37°C incubator (Firlabo, BIO BCS245).

The glass bottles containing the embryos were continuously exposed to a flow of oxygenating gas provided by a gas mixer system at a pressure of approximately 6–7 psi. The mixed gas flows from the mixer (outside the incubator) through an inlet into a humidifying water bottle located inside the incubator and then into the culture bottles. The speed of the gas flow was regulated using a valve on the gas mixer and monitored by observing the rate of bubbles formed in an outlet water-filled test tube inside the incubator. During the experiments, the bubble flow rate was maintained at 4–6 bubbles per second.

### Controlled oxygen experiments with limb explants

The rolling *ex utero* culture system was used for controlled oxygen perturbation experiments. For the mouse embryo limb development experiment shown in Fig S4A, the developing E12.5 mouse limb isolated as described above and placed into a glass bottle and cultured in 2 mL BGJb with 95% oxygen and cultured for 3 days. For the embryonic mouse, and *Xenopus* post-amputation experiments, the isolated hindlimbs were amputated (distal amputation) at the presumptive ankle, and the autopod was discarded. The remaining proximal part was transferred into the glass bottles (Schmizo, 9000.42) containing 2 mL of BGJb media for mice, and 2 mL of L- 15 media for *Xenopus*. The limbs were then cultured in the rolling system for 6 hours to 3 days with indicated oxygen amounts, as indicated in text. Mouse samples were cultured at 37°C with 5% CO₂, while *Xenopus* limbs were cultured at 22°C with 5% CO₂. Three limbs were pooled into each bottle containing 2 mL of media.

### *In vivo Xenopus* developing limb regeneration experiments

For *in vivo* developing limb amputations, *Xenopus* tadpoles (NF stages ∼53–54) were anesthetized with 0.01% MS222 (Sigma Aldrich, A5040) and the right hindlimb ankle was excised using a scalpel (As One, 2-5726-25). After amputation, tadpoles were allowed to recover in 0.1x MMR solution (0.1 M NaCl, 2 mM KCl, 1 mM MgSO₄, 2 mM CaCl₂, 5 mM HEPES, pH 7.8) and then cultured individually in single wells of 6-well plates (Thermo Scientific, 130184) with the water level set to half the container height. Tadpoles were maintained for 24 hours at 25°C in an incubator (Fukushima, FMU-054I) set to normoxia (20% O₂) or in an incubator (Astec, APM-30D) set to 60% O₂. For tissue collection, the cultured tadpoles were re-anesthetized with 0.01% MS222, and the amputated limb were excised using 30G injection needles (Nipro, 01-134).

### Small molecule inhibitor treatments

The following small molecules were included in mouse or *Xenopus* explant cultures in indicated experiments at the specified final concentrations: 50 µM TP0463518 (Selleckchem, S0729), 50 µM IOX2 (Selleckchem, S2919), 100 µM DMOG (MedChemExpress, HY-15893), 50 µM NIBR-LTSi, and 50 µg/mL Verteporfin (Sigma-Aldrich, SML0534). All small molecules were dissolved in DMSO (Thermo Scientific, BP231-100), and were diluted in media and maintained throughout the duration of experiments, with fresh molecules added during media changes, once every 2 days. The volume matching DMSO vehicle was added to as a (vehicle) control in respective experiments.

### Mouse explant AER induction experiments

For AER induction experiments in Fig 3, S8, and S9, a cocktail of small molecules and recombinant proteins were used with the following final concentration: 100 ng/mL BMP4 (PeproTech, 120-05ET), 100 ng/mL FGF10 (PeproTech, 100-26), 3 µM CHIR99021 (Calbiochem, 361559), and 100 ng/mL EGF (Thermo Scientific, AF- 100-15). CHIR99021 was prepared in DMSO (Thermo Scientific, BP231-100) and recombinant proteins were prepared according to manufacturer’s recommendations. The vehicle control had the volume matched DMSO and according to the vehicle of the recombinant protein batches. All molecules were diluted in media and maintained throughout the duration of the experiment, with fresh cocktail added during media changes once every 2 days.

### Oxygen sensor measurements

The Microx 4 fiber optic oxygen transmitter (PreSens, 200001505, 200001486) equipped with sensor type PSt7-10 and oxygen microsensors type PSt7-02 (PreSens, 200001523) was used to measure oxygen levels in the media or *Xenopus* husbandry tanks, reported in Fig 1E and S3. Briefly, the oxygen sensor was calibrated according to the manufacturer’s instructions. Then, the sensor was immersed in the media at the time of sample collection, or in *Xenopus* husbandry tanks. Then, the oxygen levels were recorded every 2 seconds for total of 8–10 measurements. Prior to recording oxygen levels, the sensor was allowed to settle into the media or tank water for about 1-2 minutes until the meter values were constant. The averaged value was recorded to determine the final levels of oxygen. The oxygen sensor was also used to verify oxygen concentrations in controlled oxygen experiments involving the MDCK cell line and the ex-utero rolling culture system.

### MDCK cell culture

MDCK II cells (Sigma-Aldrich, MTOX1300) were cultured in 75 cm² dishes (TPP, 90076) at 60–90% confluency in growth medium consisting of DMEM (Gibco, 10566016), 10% FBS (Gibco, F7524), and penicillin/streptomycin (BioConcept, 4-01F00-H). Cells used for the experiments were thawed from vials containing cells in growth medium supplemented with 10% DMSO (Sigma-Aldrich, D8418) kept in liquid nitrogen, and passage was kept between passage 2 and 10 since thawing. Prior to seeding, cells were rinsed with 1X DPBS (Gibco, 14190-094) and incubated with 1.5 mL of 0.25% Trypsin-EDTA (Company, 5-54F00-H) for 10–20 minutes until fully detached. The detached cells were resuspended in 10 mL of growth media, counted, and 335,000 cells were seeded into each well of a 12-well imaging plate (Cellvis, P12-1.5P). The volume of each well was completed to 2 mL.

After 24 hours in a standard incubator (Fisher scientific, 17154961), a media change was performed, and the plates were transferred to a custom-made controlled temperature and oxygen chamber (Life Imaging Services). Cells were cultured at 37°C, under a gas mixture with saturated humidity, 5% CO₂, either 21% or 10% O2, and N_2_ for 24 hours. Oxygen levels were continuously monitored using a dioxygen sensor throughout the experiment.

At the end of the experiment, cells were washed with 1X DPBS, fixed with 4% PFA (Sigma-Aldrich, 158127, dissolved in DPBS) for 10 minutes, washed three times with 1X DPBS, and stored in fresh DPBS until staining.

### Cryosectioning sample preparation and histology

Mouse and *Xenopus* limb explants, as well as intact limbs, were washed once in 1X PBS (0.7X PBS for *Xenopus*) and fixed in 4% formaldehyde (ThermoScientific, 119690010), diluted in 1X PBS, for 40 minutes at room temperature while rotating. The tissues were then washed three times in 1X PBS to remove the fixative and incubated in 15% sucrose (ThermoScientific, S-8600-65) for 2 hours at room temperature while rotating, followed by incubation in 30% sucrose overnight at 4°C. The limb tissues were positioned in optimal cutting temperature (OCT) solution (VWR, 361603E) under a stereomicroscope and snap-frozen in a bath of isopentane (ThermoScientific, 24872.298) and dry ice. Tissues were stored at −80°C until sectioning.

A slightly modified method was used for fixing and embedding *in vivo Xenopus* limb samples (Fig 5A). Briefly, excised limb regenerating developing limbs were fixed in 4% paraformaldehyde (diluted in 1X PBS) for 40 minutes at room temperature, transferred to 10% sucrose (Nacalai Tesque, 30404-45) in PBS for 2 hours at room temperature, and then to 30% sucrose overnight at 4°C. Samples were embedded in OCT compound (Sakura Finetek, 4583), snap-frozen by briefly touching the surface of liquid nitrogen, and stored at −80°C.

For cryosectioning, tissues were cut into 10 μm sections using a cryostat (Leica, CM1950) and placed on Superfrost Plus slides (HuberLab, 10.0344.01). The slides were stored at −80°C until used for staining.

### Whole mount alcian blue staining

Whole-mount Alcian Blue staining was performed as previously described, with minor modifications (*88*). Briefly, after fixation and washing with 1X PBS, individual limbs were placed in low-binding Eppendorf tubes and processed for staining. The limbs were dehydrated through sequential incubation in 25%, 50%, and 70% ethanol solutions for 20 minutes each, with rotation. Next, the limbs were incubated in Alcian Blue solution (3:1 ethanol to water; Millipore, TMS-010-C) overnight at room temperature, with rotation. On the second day, the samples were rehydrated by sequential washes in 70%, 50%, and 25% ethanol, followed by water, for 20 minutes each at room temperature, with rotation. Finally, the samples were treated with 1% KOH for 30 minutes, followed by incubation in a 1% KOH/20% glycerol solution at room temperature overnight or until the samples appeared cleared. The stained limbs were imaged using stereoscope microscope (Nikon, SMZ745T) attached to a camera (Nikon, DS-Fi3).

### Hematoxylin and Eosin Staining

Hematoxylin and Eosin (H&E) staining was performed using the Tissue-Tek Prisma automated slide stainer. Initially, cryosection slides were air-dried for 10 minutes. The slides were then rinsed with deionized (DI) water before being incubated with Hematoxylin (Biosystems, 3873.2500) for 5 minutes. After another DI water rinse, the slides were incubated with Eosin Y (Sigma, E4382) for 1 minute. Subsequently, the samples were dehydrated through sequential incubation in 76% ethanol, 96% ethanol, and 100% ethanol (twice), each for 1 minute. Finally, two washes with xylene were performed before mounting the samples with coverslips.

### Hybridization chain reaction (HCR)

HCR was performed as previously described (*89*), with minor modifications. On the first day, slides were washed in 1X PBS for five minutes, permeabilized in 70% ethanol for twenty minutes, and washed again with 1X PBS for five minutes. A hydrophobic (PAP) pen (Vector Laboratories, H-4000) was used to outline the tissues, and 200 μL of probe wash buffer (Molecular Instruments) was applied to the sections for ten minutes. After removing the wash buffer, 200 μL of hybridization buffer (Molecular Instruments) was added to the tissue, and the samples were incubated at 37°C for 30 minutes. Meanwhile, the probe solution was prepared by diluting mRNA-targeting probes in 200 μL of hybridization buffer to a final concentration of 30–40 nM. This solution was also incubated at 37°C for 30 minutes. After removing the hybridization buffer from the samples, the probe solution was applied, and the samples were incubated for 12–16 hours at 37°C. On the second day, samples were washed twice for ten minutes with wash buffer and twice for twenty minutes with 5X SSC-T at room temperature. During the 5X SSC-T washes, the amplification solution was prepared. The fluorophore-tagged hairpins (h1 and h2) corresponding to the probes were first heated to 95°C for 90 seconds. The hairpins were then allowed to return to room temperature in the dark for 30 minutes before being added to the amplification buffer (Molecular Instruments) at a final concentration of 40–60 nM in 200 μL. Amplification buffer without hairpins was applied to the samples for 10 minutes, followed by the final amplification solution containing hairpins, which was incubated at room temperature, protected from light, for 12–16 hours. On the third day, samples were washed twice for twenty minutes with 5X SSC-T and mounted using DAPI+ mounting media (Vector Laboratories, H-1200) and coverslips (Sigma-Aldrich, BR470050). HCR probes (*fgf8.L* (*Xenopus*), *Fgf8* (Mouse)) and hairpins were purchased from Molecular Instruments.

### Immunostaining

Slides containing cryosectioned samples were washed three times with 1X PBS for five minutes, followed by three washes with PBS-T (1X PBS supplemented with 0.1% Triton). Please note that for HIF-1 alpha antibody slides were incubated in cold methanol at −20°C for 10 minutes followed by another three 1XPBS and 1X PBST washes, as reported before (*90*). The samples were then incubated in a blocking solution consisting of 50% Cas-Block (Thermo, 008120) and 50% PBS-T for one hour at room temperature. After removing the blocking solution, primary antibodies diluted in the same blocking solution were applied to the samples and incubated overnight at 4°C. The following day, the primary antibody solution was removed, and the slides were washed three times with PBS-T for five minutes each. Secondary antibodies, specific to the host species of the primary antibodies, were diluted in blocking solution and applied to the tissue for one hour at room temperature in the dark. After incubation with the secondary antibodies, the samples were washed three times with PBS-T for ten minutes each, followed by three additional ten-minute washes with 1X PBS. Finally, the samples were mounted using DAPI+ mounting media and coverslips.

For MDCK cell line staining cells were washed 3 times with 1X PBS for five minutes, followed by three washes with PBS-T. The cells were then incubated in Phalloidin (1:6000) and Hoechst (1:5000) diluted in PBS-T for 1hr. After incubation with staining solution, cells were washed 3 times with 1X PBS for five minutes, followed by three washes with PBS-T and stored at 4 degrees until imaging.

### Antibodies and working dilutions

Primary antibodies and used dilutions were as follows: HIF-1 alpha (D1S7W) XP (Cell Signaling, 36169S, 1:200), Anti-active YAP1 antibody (Abcam, 205270, 1:200), Phalloidin-iFluor 488 Reagent (Abcam, 176753, 1:6000), TP63 [4A4] (Abcam, ab735, 1:200), and Lamin A/C (4C11) (Cell Signaling, 4777, 1:100). Secondary antibodies were as follows: AlexaFluoro 488 Goat anti-Rabbit (Invitrogen, A32731, 1:400), AlexaFluoro 647 Donkey anti-Rabbit (Invitrogen, A31573, 1:400), AlexaFluoro 488 Donkey anti-Mouse IgG (H+L) (Invitrogen, A21202, 1:400), or AlexaFluoro 594 Donkey anti-Mouse (Invitrogen, A21203, 1:400).

### Confocal and brightfield imaging

All H&E images were aquired using Olympus VS200 UPLXAPO with 40x/0.95 objective. Confocal images in Fig 1G, Fig 2H (left), Fig 3A, Fig S4B & D, Fig S6E were taken with a Leica SP8 upright confocal microscope with 20x/0.75 HC PL APO. Confocal images in Fig 2A-E, G & H (right), Fig 5 C, E, I, Fig S5A & C, Fig S6A, C, D, and F were taken either with an Upright Leica DM6 CS or Inverted Leica SP8 FLIM confocal microscope with 63x/1.40 and 63x/1.30 HC PL APO objectives respectively. Confocal image in Fig 2J was taken with Inverted Leica DMi8 confocal microscope with 63x/1.40 HC PL APO objectives. All images were analyzed using Fiji. Optical-sections, or max-projection of z-stacks were used and Fiji was used to adjust the contrast to emphasize biological relevance. When necessary, images were cropped, flipped, and/or rotated to highlight biological relevance.

### Confocal image analysis

For phalloidin-based measurements (Figures 2, 5, S5 and S6), cells that were either TP63+ or at the top of the basement membrane were considered basal epithelial cells and manual quantification via Fiji was used. The cell elongation ratio was measured manually in Fiji by taking the ratio of the width and height of individual cells with respect to the basement membrane. The first most distal twenty cells were counted for these measurements. Apical-basal actin ratio was measured by taking the ratio of apical to basal actin intensity across a group of basal epithelial cells in each section. For basal actin thickness, the most representative region was determined visually and quantified for each section.

For both YAP and LAMINA/C quantification (Figs 2, 5, S5 and S6), cells were classified as YAP or LAMINA/C positive if they showed nuclear signals covering more than 50% of the nuclear area after contrast adjustment of the images. Cells with no detectable nuclear signal or faint puncta were classified as YAP or LAMINA/C negative. The first most distal twenty cells were counted for LAMINA/C measurements.

For MDCK cell line-based quantifications (Fig S7), aspect ratio, number of neighbours per cell (determined by identifying neighbouring cells in contact along their phalloidin-stained borders) and nuclear area were measured manually. Crowding was estimated by calculating the ratio of nuclei counted to the area of each image.

### Sample preparation for single-cell mRNA sequencing (scRNA-Seq)

For mouse scRNA-seq experiments, 5 limb explants for 3 dpa Fig 3B, and S8, and 6 limb explants for 1 dpa Fig 4H, and S14 were pooled in an Eppendorf tube and immediately processed for dissociation. The tissues were first washed with 1X PBS and then treated with 500 µL of 0.25% trypsin-EDTA (Gibco, 25200056) at 37°C for 5 minutes. Cells were mechanically dissociated by gently pipetting 5–8 times until they were visibly dislodged. To halt the enzymatic reaction, 1 mL of DMEM (Gibco, 61965-026) supplemented with 10% FBS (Gibco, 16141- 079) was added. The cells were centrifuged at 600 rcf for 5 minutes at room temperature and subsequently washed with 1 mL of 1X PBS containing 0.04% BSA (Gibco, 15260037). Finally, the cells were passed through a 40 μm cell strainer (Sigma-Aldrich, BAH136800040) and counted manually using trypan blue and with the Countess 3 automated cell counter (Thermo Scientific, AMQAX2000).

For *Xenopus* scRNA-seq (Fig 5J and S19), limb explants from eight embryos were pooled in an Eppendorf tube and immediately processed for dissociation, as described previously (*11*). The tissues were washed with 0.7X PBS and treated with 0.35 mg/mL Liberase solution (Sigma-Aldrich, 05401127001) diluted in 0.7X PBS for 30 minutes at room temperature while gently rotating. Cells were mechanically dissociated by pipetting 10–15 times until visibly dislodged. Next, 1 mL of L15 medium (supplemented with 1% antibiotic-antimycotic, 0.8 mM L- glutamine, and 20% FBS) was added to the mixture. The cells were then centrifuged at 600 rcf for 5 minutes at room temperature and washed with 1 mL of 0.7X PBS containing 0.04% BSA. Finally, the cells were passed through a 40 μm cell strainer and counted manually using trypan blue and with the Countess 3 automated cell counter.

After isolation, mouse cells were washed once with PBS, 0.04% BSA, while *Xenopus* cells were washed once with 0.7x PBS, 0.04% BSA. After filtration through a 40 um Flowmi strainer (Bel-Art, Wayne, NJ), cells were resuspended in PBS 0.04% BSA for mouse cells and 0.7x PBS 0.04% BSA for *Xenopus* cells. All cells were checked for absence of significant doublets or aggregates, and loaded into a Chromium Single Cell Controller (10x Genomics, Pleasanton, CA) in a chip together with beads, master mix reagents (containing RT enzyme and poly-dt RT primers) and oil to generate single-cell-containing droplets. For mouse samples, 4’500 cells were targeted, for *Xenopus* samples, 10’000 cells were targeted.

Single-cell Gene Expression libraries were then prepared using the Chromium Single Cell 3’ Library & Gel Bead Kit v3.1 (PN-1000268) following the manufacturer’s instruction (protocol CG000315 Rev C). Quality control was performed with a TapeStation 4200 (Agilent) and QuBit dsDNA high sensitivity assay (Thermo) following manufacturer instructions. The sequencing libraries were loaded on NovaSeq 6000 (Illumina) and/or Aviti Cloudbreak Freestyle (Element Biosciences) flow cells and sequenced according to manufacturer instructions, with a read1 of 28nt and a read2 of 90nt, at a depth of ca 50-95k reads/cell. BCLconvert v 3.9.3 (Illumina) and bases2fastq v1.8.0 (Element Biosciences) were used to demultiplex reads, which were then quality-controlled with fastQC v0.12.1. FastQ Screen v0.16.0 tool was used for screening FASTQ files reads against multiple reference genomes.

### Sample preparation for single-nuclei mRNA and ATAC (scMultiome) sequencing

For scMultiomics (Fig 4, and S10), four intact E12.5 limbs and eight cultured limb explants from different time points (6, 24, 72 hpa) were collected. The tissues were pooled into an Eppendorf tube, snap-frozen on dry ice, and stored at −80°C until use. Single nuclei dissociation was conducted using a modified version of the 10x Genomics protocol (CG000365 Rev B). Briefly, the tissues were treated with 0.05X lysis buffer (50 μL) and mechanically dissociated using an electric pellet pestle (Kimble, 749540-0000) applying 2–5 short pulses. Wash buffer (150 μL) was then added to the suspension, and the mixture was further dissociated by pipetting up and down 10 times. The resulting suspension was passed through a 40 μm cell strainer and centrifuged at 600 × g for 5 minutes at 4°C using a swing-bucket centrifuge. The nuclei pellet was washed, centrifuged two additional times, and resuspended in 1X nuclei dilution buffer. Finally, the nuclei suspension was passed through a 40 μm Flowmi Cell Strainer for further refinement. Detailed buffer recipes, reagent information, and the complete protocol are available in the 10x Genomics Protocol (CG000365 Rev B).

After isolation, mouse nuclei were filtered through a 40 um Flowmi strainer (Bel-Art, Wayne, NJ) and resuspended in Nuclei Diluted Buffer, prepared according to the manufacturer’s instruction (protocol CG000338 Rev F). and filtered once more through a 40 um Flowmi strainer. All nuclei were checked for absence of significant doublets or aggregates and incubated in a Transposition Mix. After transposition, nuclei were loaded into a Chromium Single Cell Controller (10x Genomics, Pleasanton, CA) in a chip together with beads, master mix reagents (containing RT enzyme, poly-dt RT primers and a Spacer sequence that enables barcode attachment to transposed DNA fragments) and oil to generate single-cell-containing droplets. For these samples, 3’000 or 6000 nuclei were targeted.

Single-cell Gene Expression libraries and ATAC libraries were then prepared using the Chromium Next GEM Single Cell Multiome ATAC + Gene Expression v1.0 (PN- 1000285) following the manufacturer’s instruction (protocol CG000338 Rev F). Quality control was performed with a TapeStation 4200 (Agilent) and QuBit dsDNA high sensitivity assay (Thermo) following manufacturer instructions. The sequencing libraries were loaded on NovaSeq 6000 (Illumina) flow cells and sequenced according to manufacturer instructions. For Gene expression libraries read1 was 28nt and read2 was 90nt, and the sequencing depth was about 75-105k reads/cell. For ATAC libraries read1 and read2 were 49nt, and the sequencing depth was about 85-175k reads/cell. BCLconvert v 3.9.3 (Illumina) was used to demultiplex reads, which were then quality-controlled with fastQC v0.11.9. FastQ Screen v0.14.0 tool was used for screening FASTQ files reads against multiple reference genomes.

### Single nuclei mRNA and ATAC (scMultiome) analyses

Raw FASTQ files from both RNA and ATAC sequencing were processed using Cell Ranger ARC (version 2.0.2) with the mm10 mouse genome reference dataset provided by 10x Genomics (*cellranger-arc-mm10-2020-A- 2.0.0*). The processed data were further analyzed using in-house R-scripts and R (v4.3.1) with packages Seurat (v4.3.1) and Signac (v1.10).

Seurat objects for each dataset were created using the RNA assay, importing pre-filtered raw count matrices (filtered_feature_bc_matrix.h5) with Read10X_h5 and CreateSeuratObject, and further filtering cells based on a minimum of 5 cells and 200 features. The objects were merged using *merge* method and processed through standard steps: data normalization (LogNormalize method, scaling by 10,000 and log-transforming), scaling to a mean of 0 and variance of 1, and PCA on the top 2,000 variable genes (identified by VST), retaining the first 30 components. Neighboring cells were identified and clustered with FindClusters using the Louvain algorithm (resolution = 0.8), followed by UMAP visualization based on the first 30 PCA components.

Cell type annotation was performed manually using the expression of marker genes with previously reported genes (*24*). CT subpopulations were defined accordingly to their clusters of origins, without using FindSubCluster. CT *Sox9^Low^* and CT Sox9*^High^*subpopulations were identified based on *Sox9* gene expression in CT population, with a threshold set at 0.5.

The ATAC-seq assay was added to the merged Seurat object following a standard integration pipeline. First, a unified peak set was created by merging overlapping peaks from each dataset using the reduce method in the GenomicRanges. Peaks were called using the Signac function CallPeaks integrating macs3, followed by removal of non-standard chromosomes and peaks in blacklist regions with functions keepStandardChromosomes and subsetByOverlaps. A chromatin assay was then created by constructing a feature matrix with the FeatureMatrix function, using the fragment files and the unified set of peaks, followed by CreateChromatinAssay. The ATAC-seq assay was processed according to Signac’s recommendations, including normalization and variable feature identification with RunTFID (Term Frequency-Inverse Document Frequency) to account for varying sequencing depths across single cells and FindTopFeatures. Finally, a joint UMAP was generated using the weighted shared nearest neighbor (wsnn) method, incorporating the first 50 principal components (PCs) of the RNA-seq data and the first 50 LSI components of the ATAC-seq data through FindMultiModalNeighbors.

A similar process was applied to identify chromatin accessibility peaks at each time-point and for each cell-type (peak calling with CallPeaks followed by the chromosome and blacklist filtering step).

### snATAC-Seq peak annotation and differentiation chromatin accessibility

Peaks were annotated with annotatePeak (R package ChIPseeker (v1.24.0))), using library TxDb.Mmusculus.UCSC.mm10.knownGene, defining TSS regions as TSS +/-1kb and keeping the default genomic annotation priority.

Differential chromatin accessibility was assessed between 6 hpa and intact E12.5 using FindMarkers with the LR test on peaks called for each cell type and condition as described above, with a minimum of 0.1% expression, a log fold-change threshold of 0.25, and at least 5 cells per feature and group. Up to 150 cells per condition were included, with nCount_peaks used as a latent variable.

Coverage plots were generated using the Signac method CoveragePlot, which displays normalized signals obtained by calculating the averaged frequency of sequenced DNA fragments for the different groups of cells considered.

The heatmap of the AER gene set chromatin accessibility at each time point in basal epithelial cells were generated using the EnrichedHeatmap package (v1.29.3). Matrices used were computed from individual ATAC- seq signal tracks derived from basal epithelial cells at each time point, using normalizeToMatrix with settings for signal normalization, smoothing, background correction, and value scaling. These signal tracks were prepared using an adapted version of the Signac function ExportGroupBW (https://github.com/stuart-lab/signac/blob/e65a74dbe8e489f92b01a2a9c049c67a317c83a0/R/utilities.R), with 50 bp tile sizes and normalization of fragments per tile to the number of cells.

### Single cell mRNA Sequencing (scRNA-Seq) analyses

Mouse and *Xenopus* scRNA-seq samples were processed using Cell Ranger (version 8.0.1) with the GRCm39 (2024-A) mouse genome reference dataset provided by 10x Genomics Genomics (*refdata-gex-GRCm39-2024- A*) and the *Xenopus laevis* 10.1 (*CRREF007_Xenopus_Laevis_v10.1_Genome*), for *Xenopus* samples. The number of expected cells was set to 4500, and intronic reads were retained. The remaining analysis were mostly performed within R (v4.3.1) using in-house R-scripts and R packages Seurat (v4.3.1) and Signac (v1.10). The count matrices were imported into Seurat. Cells were filtered to exclude those with fewer than 500 or more than 6,000 detected genes, more than 5% mitochondrial UMI counts, or UMI counts outside the range of 5000 to 30,000. Data were normalized using the LogNormalize method (scaling by 10,000 and log-transforming counts), then scaled to a mean of 0 and variance of 1. Principal component analysis (PCA) was performed on the top 2,000 variable genes (identified by VST), retaining the first 30 components. Neighboring cells were identified using these 30 components, and clustering was performed using the Louvain algorithm at a resolution of 0.8. UMAP was then applied to the first 30 PCA components for visualization.

### ScRNA-seq data integration and downstream analyses

To address the potential absence of certain cell-types in some datasets, we performed RPCA-based integrations as described in (https://satijalab.org/seurat/archive/v4.3/integration_rpca.html). Specifically, we used the SelectIntegrationFeatures method with 3,000 features, followed by PCA on these selected features. Anchors were then identified using the FindIntegrationAnchors method, based on the first 20 principal components from the RPCA reduction. using the IntegrateData function, with integration performed on the first 20 principal components.

Cell cycle correction was performed as described in (*24*) following the sample integration. Briefly, cell cycle scores were assigned using the CellCycleScoring method with a list of cell-cycle genes from Seurat (updated with 2019 gene symbols). For *Xenopus* samples, we added .S and .L to the Mouse gene symbols. Principal components (PCs) with summed loading values for these genes exceeding sample-specific thresholds were classified as cell cycle-correlated. PCA was then re-run, excluding the top 10% of genes associated with these cell cycle-correlated PCs.

PCA and cell clustering of the integrated object were done as for individual datasets. Cell type annotation was performed manually using the expression of marker genes with previously reported genes (*24*), using the clusters obtained at a resolution of 0.8 (integrated_snn_res.0.8). CT subpopulations were defined accordingly to their clusters of origins, without using FindSubCluster. *Sox9^Low^* CT and *Sox9^High^* CT subpopulations were distinguished within the CT population based on *Sox9* gene expression levels, with a threshold of 0.5 unless otherwise stated.

Differential expressions between samples and/or cell-types were assessed using the FindMarkers function in Seurat with the Wilcoxon rank sum test. Genes with a log-fold change > 0.1 and expressed in at least 1% of cells were considered, including both up- and down-regulated genes.

Functional analysis was performed using gost method (gprofiler2 package) on both up- and down-regulated genes (adjusted p-value < 0.05), with a significance threshold of 0.05 and default settings for other parameters. For *Xenopus*, gene names were converted to mouse format by removing the .S and .L.

Gene set enrichment scores were calculated using AUCell, as before (*24*), with default settings for the following genes sets (**Supplemental Table 4**).

Mouse, *Xenopus*, human, axolotl – publicly available cross-species limb development and regeneration datasets (Fig. 6B and Fig. S21A) were obtained and processed as previously described (*24*).

### Sample preparation for CUT&Tag sequencing

For CUT&Tag (Fig S15, S17, and S18), single-cell suspensions were prepared as described above for single-cell mRNA sequencing. For mouse samples, five limb explants were pooled together, and for *Xenopus*, eight amputated limbs were pooled together.

CUT&Tag libraries were prepared as previously reported (*93*). The centrifuged cells were resuspended in 1 ml of CUT&Tag wash buffer (20 mM HEPES [pH 7.5] (Jena Bioscience, BU-106-75), 150 mM NaCl (Sigma-Aldrich, 71386), 0.5 mM spermidine (Merck, S2626), 5 mM sodium butyrate (Sigma-Aldrich, 303410), and 1X Protease Inhibitor Cocktail (Roche, 11873580001) in UltraPure distilled water (Invitrogen, 10977-035)). Then, a volume of Concanavalin A beads (Polysciences, 86057-3) corresponding to 15 µl per histone mark target was prepared by removing the supernatant on a magnetic rack and washing in 1 ml binding buffer (20 mM HEPES [pH 7.5], 10 mM KCl (Sigma-Aldrich, 60142), 1 mM CaCl_2_ (Sigma-Aldrich, 21115), and 1 mM MnCl_2_ (Sigma-Aldrich, M1028) in UltraPure distilled water). The supernatant was removed again, and the beads were resuspended in the original volume (15 µl per sample) of binding buffer.

Beads were added to the cell suspension for binding. Using all cells from the dissociated limbs, this corresponded to 60k-600k live cells per histone mark. After 15 minutes of incubation at room temperature on a rotating wheel, tubes were placed in a magnetic rack, the supernatant was discarded, and cells were permeabilized in wash buffer supplemented with 0.01% Digitonin (Invitrogen, BN2006) until nuclei were visible around the beads via Trypan Blue (Invitrogen, T10282) staining. The supernatant was discarded again on the magnetic rack, and samples were resuspended in 150 µl wash buffer per histone mark and distributed in separate tubes for antibody binding. Primary antibodies were added as per the manufacturer’s suggested concentration for CUT&Tag (H3K27me3 (Diagenode, C15410195): 1 µg in 150 µl; H3K4me3 (Diagenode, C1541003): 0.5 µg in 150 µl; H3: dilute at 1:150 (Activemotif, 39763) along with 2 mM EDTA (Invitrogen, 15575- 038). Samples were incubated overnight at 4 °C on a nutator. The next day, the primary antibody solution was removed on a magnetic rack, and the beads were washed twice with 400 µl of CUT&Tag wash buffer, vortexing briefly each time. Beads were resuspended in 150 µl of CUT&Tag wash buffer with 1.5 µl of secondary antibody (Abcam, 46540; Antibodies Online, ABIN101961), and incubated for 1 hour at 4 °C on a nutator.

Next, the secondary antibody solution was removed on a magnetic rack, and the beads were washed twice with 400 µl of CUT&Tag wash buffer, vortexing briefly each time. Beads were resuspended in 150 µl of high salt wash buffer (20 mM HEPES [pH 7.5], 300 mM NaCl, 0.5 mM spermidine, 5 mM sodium butyrate, and 1X Protease Inhibitor Cocktail in UltraPure distilled water) with 2 µl of homemade Protein A-Tn5 (*94*), and incubated for 1 hour at room temperature in a thermomixer at 300 rpm. The Protein A-Tn5 solution was removed on a magnetic rack, and the beads were washed twice with 400 µl of high salt wash buffer. Beads were resuspended in 100 µl CUT&Tag wash buffer supplemented with 10 mM MgCl_2_ (Invitrogen, AM9530G) for chromatin digestion for 1 hour at 37 °C in a thermomixer at 300 rpm. The reaction was stopped by adding 4.5 µl of 0.5 M EDTA, 2.76 µl of 20% SDS (Sigma-Aldrich, 05030), and 1 µl of 20 mg/ml Proteinase K (Invitrogen, AM2546), and incubating in a thermomixer at 300 rpm for 30 minutes at 55 °C, followed by 20 minutes at 70 °C.Each sample was then purified using the ChIP DNA Clean & Concentrator kit (Zymo, D5205) as per the manufacturer’s instructions for a 100 µl sample. At the final step, samples were eluted in 25 µl of elution buffer (Qiagen, 19086) containing 2 pg of lambda-DNA (NEB, N3011S; tagmented with Tn5 and cleaned up on columns before use) as a spike-in.

For library preparation, 21 µl of each sample was transferred to PCR tube strips, and 25 µl of Nebnext® High-Fidelity 2X PCR Master Mix (NEB, M0541S) was added, along with 2 µl of each i7 and i5 index primer (*93*). The following thermocycler program was used: 1 5 min at 58 °C (gap filling); 1x 5 min at 72 °C (gap filling), 1x 30 s at 98 °C (denaturation), 14x 10 s at 98 °C (denaturation) plus 30 s at 63 °C (annealing/extension), 1x 1 min at 72 °C (extension), hold 4 °C. A final clean-up was performed by mixing in 45-55 µl of AMPure XP beads (Beckman Coulter, A63881) and incubating at room temperature for 15 minutes. The supernatant was removed on a magnetic rack, and the beads were washed twice in 400 µl 80% EtOH. Finally, beads were resuspended in 33 µl 0.1X TE buffer and incubated for 5 minutes at room temperature. On a magnetic rack, 28 µl of the supernatant was transferred to a new tube. Quality control was performed using the Qubit dsDNA HS Assay kit (Invitrogen, Q32854) and the Agilent High Sensitivity D5000 ScreenTape system (ScreenTape: Agilent, 5067- 5592; Reagents: Agilent, 5067-5593). An equimolar pool (5-15 nM) of samples were sequenced.

CUT&Tag libraries, bearing combinatorial dual indexes, were loaded at 7 or 9 pM on an Aviti Cloudbreak Freestyle flow cell (Element Biosciences) and sequenced according to manufacturer instructions, yielding pairs of 75 nucleotides reads at a depth of about 20 mio reads pairs per sample. Reads were trimmed of their adapters with bases2fastq version 1.8.0 (Element Biosciences) and quality-controlled with fastQC v0.12.1. FastQ Screen v0.16.0 tool was used for screening FASTQ files reads against multiple reference genomes.

### CUT&Tag pre-processing, alignment, spike-in normalization and combining replicates

Two sets of hybrid genomes consisting of the mouse genome (mm39, release 110: https://ftp.ensembl.org/pub/release-110/fasta/mus_musculus/dna/) with Escherichia phage Lambda (https://www.ncbi.nlm.nih.gov/datasets/genome/GCF_000840245.1/), and *Xenopus laevis* (v10.1: https://www.xenbase.org/xenbase/static-xenbase/ftpDatafiles.jsp) with the same Escherichia phage Lambda were built using the createIndices pipeline from snakePipes (*95*) (version 2.8.0).

Depending on the species, CUT&Tag sequencing data was aligned to the respective hybrid genome using the DNAmapping pipeline from snakePipes with Bowtie2 aligner and default parameters. The resulting alignment BAM files were converted to bigWig files using bamCoverage from deepTools (*96*) (version 3.3.0) and subsequently scaled based on the number of reads mapped to the spike-in regions. Spike-in scaled bigWig files were log2 normalized with wiggletools (*97*) (offset 1).

Principal component analysis was performed by calculating the average scores for each spike-in and log2 normalized bigWig files on 10000 bp genomic regions using multiBigwigSummary. For replicates with high Spearman’s correlation, the mean was calculated using wiggletools.

### CUT&Tag peak calling

Broad peaks were called from the BAM files of each sequencing run using MACS3 (*98*) (v3.0.1) with the parameters --broad-cutoff 0.1, --nolambda and the corresponding H3 BAM file (same day and same culture condition) as control. The fraction of reads in peaks (FRiP score) was calculated for each BAM file using featureCounts (*99*) from subread (v2.0.2), comparing the reads to the corresponding broad peaks.

### CUT&Tag differential analysis of different culturing conditions

To calculate enrichment scores for regions of interest (AER, CT blastema and positional information, **Supplemental Table 4**), normalized bigWig files were summarized over the regions specified in the BED files using bedtools (*100*) map (v2.30.0). The resulting bigWig signals on regions were used to compute the log2 fold change between 96-well and ALI samples for each replicate.

In addition to the regions of interest, TSS was used to quantify the global histone profiles. For visualization, heatmaps were generated on the regions of interest using computeMatrix scale-regions with parameters -- binSize 100, --missingDataAsZero and then plotHeatmap from deepTools.

### Statistics and replicate information

Throughout this manuscript, *n* indicates technical replicates (e.g., different animals, cell culture wells, cells), while *N* indicates independent biological replicates performed on separate days, using specimens from different batches or matings, or both. The number of replicates for each experiment is provided in the figure legends. Sample sizes were not predetermined, and all analyses were conducted without blinding. Unless otherwise stated, data are presented as mean ± SEM. For statistical comparisons, we used t-test for pairwise comparisons, and one-way ANOVA for comparisons involving more than two conditions. Significance levels throughout this work are represented as *p<0.05, **p<0.001, ***p<0.0001, and ****p<0.00001 unless otherwise stated.

## Supplementary tables

Supplemental_Table_1_scRNA-Seq_population

Supplemental_Table_2_DEallGenes_AllCells_Xen_96w_vs_Xen_ALI_1dpa

Supplemental_Table_3_DEallGenes_AllCells_Mouse_96w_vs_Mouse_ALI_1dpa

Supplemental_Table_4_genes_for_AUCell_Cut&Tag

---

**Figure S1.**
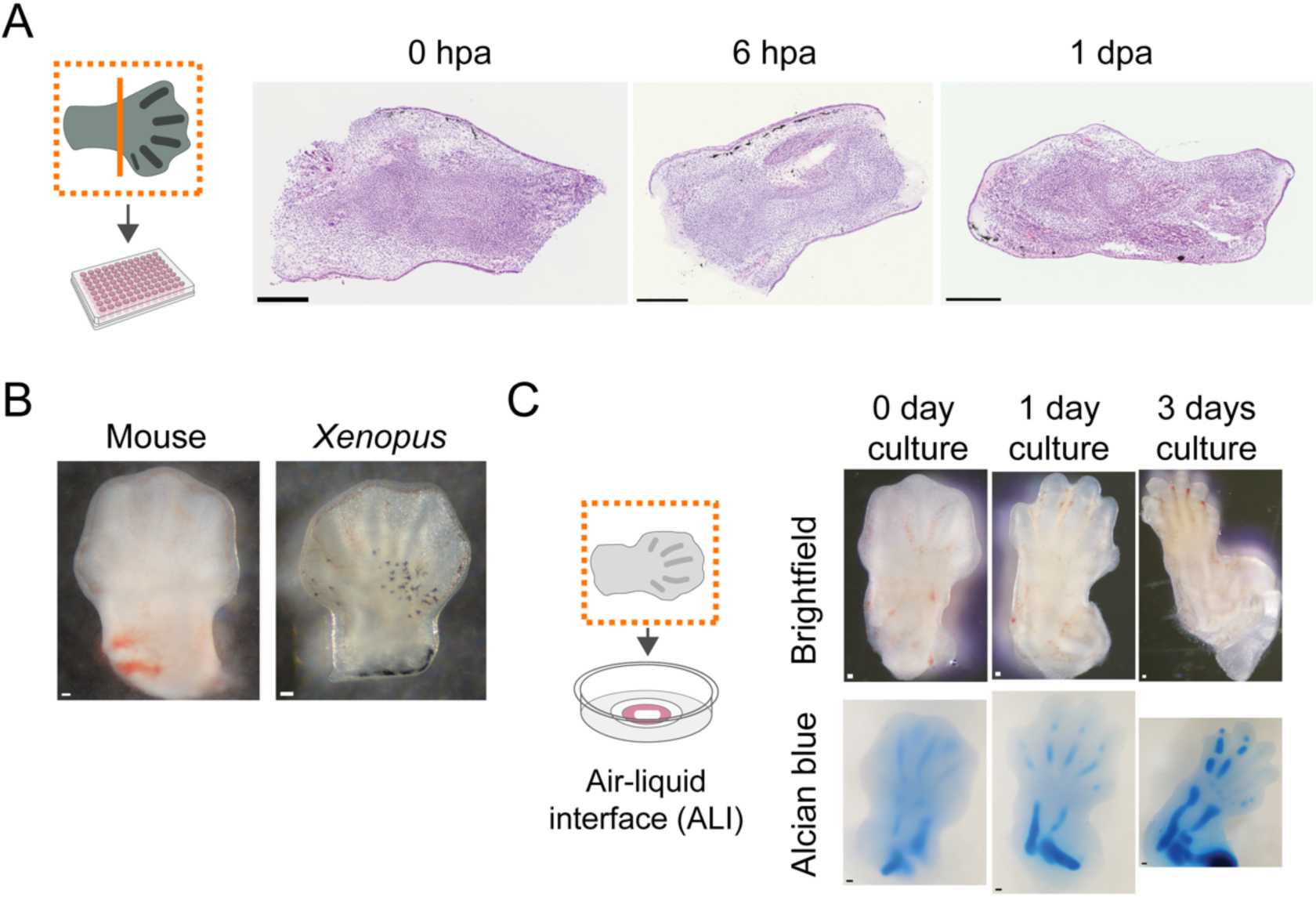
Time-course images of *Xenopus* regenerating limb explants and mouse limb development in ALI cultures. **(A)** Representative time-course histology images of *Xenopus* regenerating limb explants over the indicated time points. Scale bar = 200 μm. **(B)** Images comparing *Xenopus* NF Stage 54 and mouse E12.5 limbs, highlighting their morphological similarities and representing samples used in this study. Scale bar = 100 μm. **(C)** Representative brightfield and whole-mount alcian blue images of mouse developing limb explants, as described previously (*25*). Scale bar = 100 μm.

**Figure S2.**
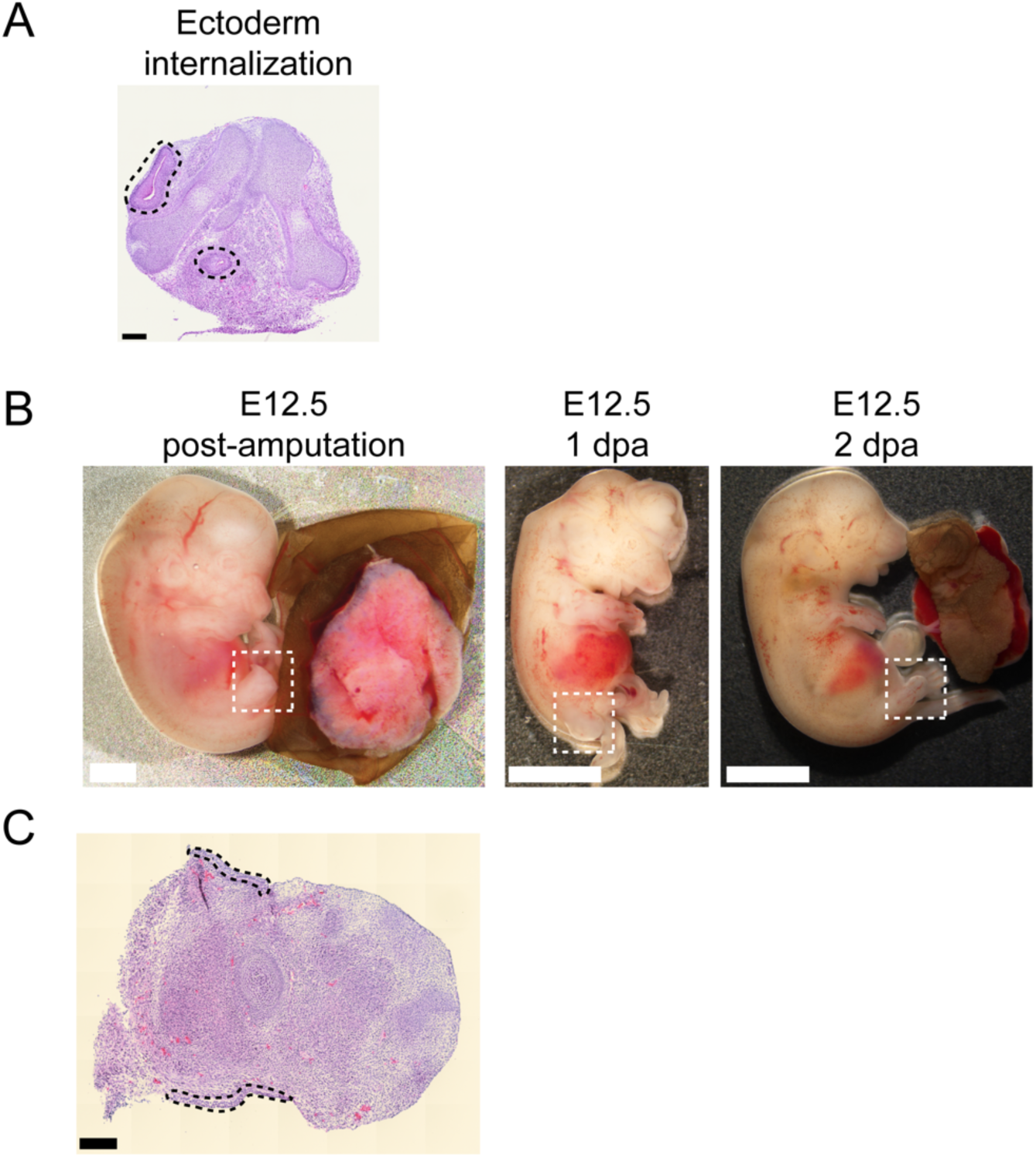
Additional images of mouse explants and whole embryos in *ex utero* culture for 1 and 2 days. **(A)** Example image of epithelial cell internalization in mouse explants cultured in ALI. Dashed lines surround the internalized epithelial cells. Scale bar = 200 μm. **(B)** Brightfield images of *ex utero* cultured whole E12.5 mouse embryos right after hindlimb amputation, 1 day, and 2 days of culturing. The embryos show signs of sub-optimal development as evidenced by e.g., their smaller size and malformed head development. Dashed boxes show amputated hindlimbs and their growth over time. Scale bar = 1000 μm. **(C)** Representative histology image of an amputated mouse hindlimb from *ex utero* cultured E12.5 mouse embryo for 1 day. Dashed lines surround the epithelial cells. Scale bar = 200 μm. N = 2, n = 0/8 embryos closed distal wound area.

**Figure S3.**
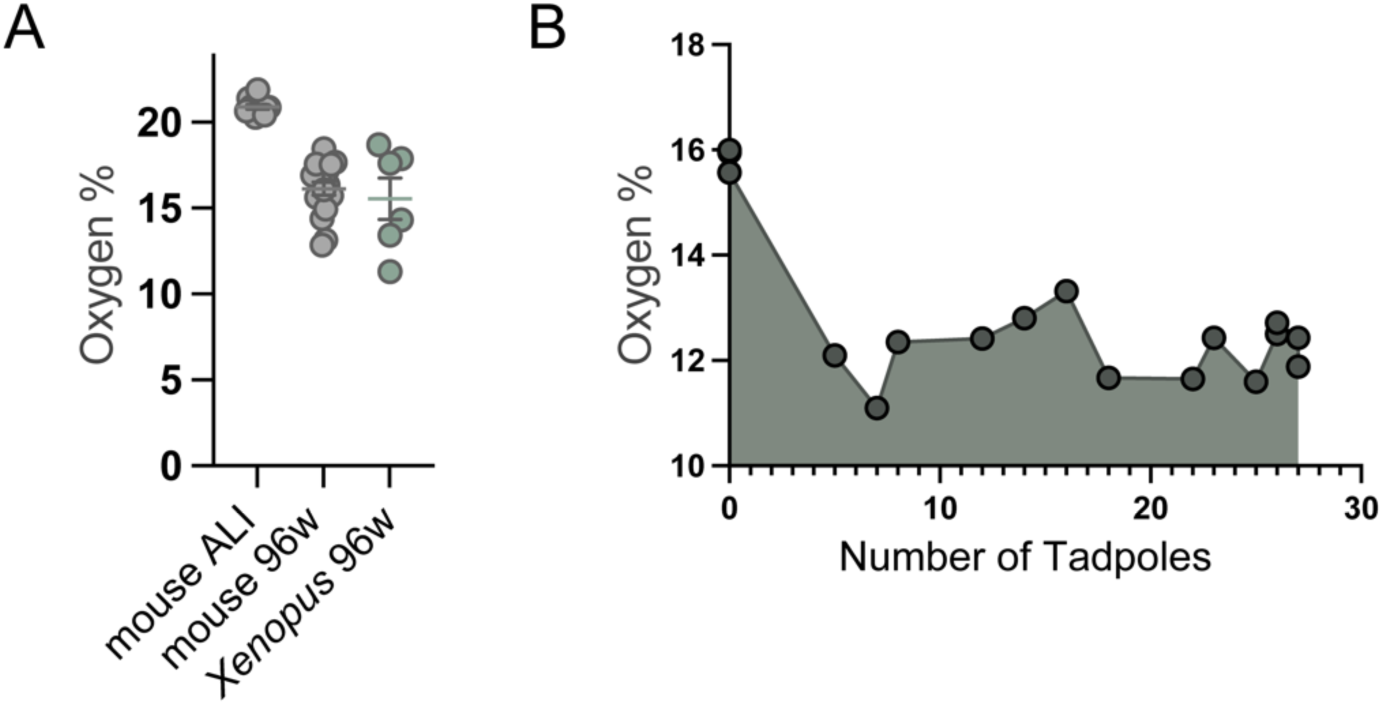
Fiber-optic oxygen sensor measurements in culture conditions and *Xenopus* husbandry tanks. **(A)** Oxygen levels measured in mouse ALI cultures, mouse 96w, and *Xenopus* 96w after 3 days of culture (*1 day after media change*). Mouse ALI N = 3, n = 11; Mouse 96w N = 3, n = 17; Xenopus 96w N = 2, n=6. Please note that this panel includes data from Figure 1E for mouse ALI and 96w, supplemented here for comparison with *Xenopus* 96w. **(B)** Oxygen measurements in *Xenopus* tadpole husbandry tanks at various time points following 4 days after weekly husbandry water change. The X-axis represents the number of animals, and the Y-axis indicates the measured oxygen levels. 14 tanks were recorded with tadpole numbers ranging from 0 to 27 as shown in the figure.

**Figure S4.**
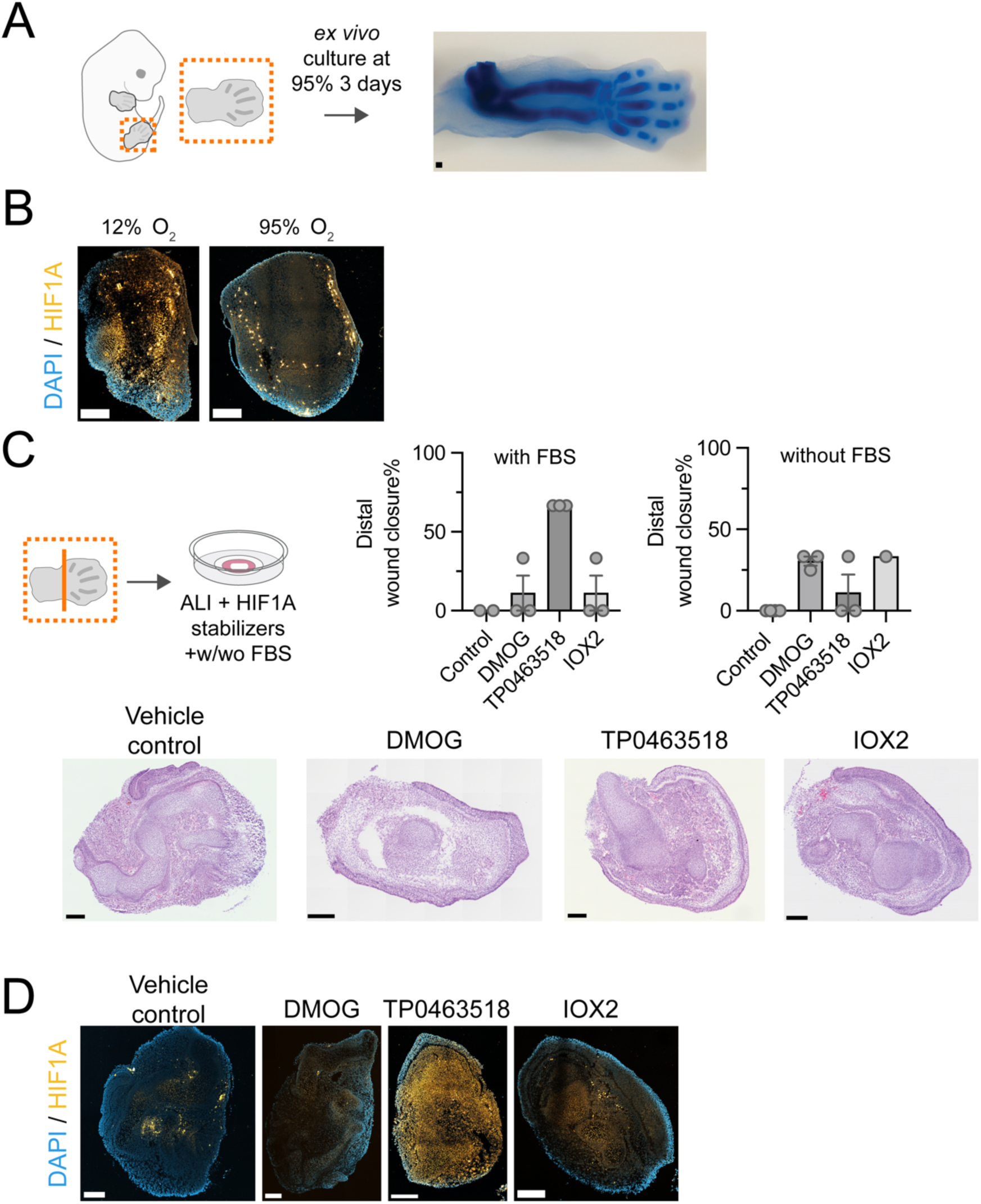
High oxygen does not impair *ex vivo* limb development, and HIF1A activation is observed under low oxygen or HIF1A-stabilizing treatments. **(A)** Representative whole-mount alcian blue staining of E12.5 mouse limbs cultured at 95% oxygen for 3 days, showing *in vivo*-like limb morphogenesis. N = 2, total n = 6. **(B)** Confocal sections of HIF1A immunofluorescence in mouse explants cultured at 12% oxygen or 95% oxygen for 6 hours. Scale bar = 250 μm. N = 2 n = 6. **(C)** Treatment of HIF1A stabilizers and distal wound healing response with or without FBS (serum) addition. Please note the TP0463518 data without serum is also shown in Fig 1H. Representative histology images of mouse explants cultured in ALI with HIF1A-stabilizing molecules. Each dot represents individual sections. Scale bar = 200 μm. With serum: Vehicle control N=2, n=0/9 explants closed distal wound area; DMOG N=3, n=1/9 explants closed distal wound area; TP0463518 N=3, n=6/9 explants closed distal wound area; IOX2 N=3, n=1/ explants closed distal wound area. Without serum: Vehicle control N=3, n=0/9 explants closed distal wound area; DMOG N=3, n=3/10 explants closed distal wound area; TP0463518 N=3, n=1/9 explants closed distal wound area; IOX2 N=1, n=1/3 explants closed distal wound area. **(D)** Confocal images of HIF1A immunofluorescence in mouse explants cultured in ALI with HIF1A-stabilizing molecules for 3 days. Vehicle treatment serves as the control. Scale bar = 250 μm. N = 2, n = 6.

**Figure S5.**
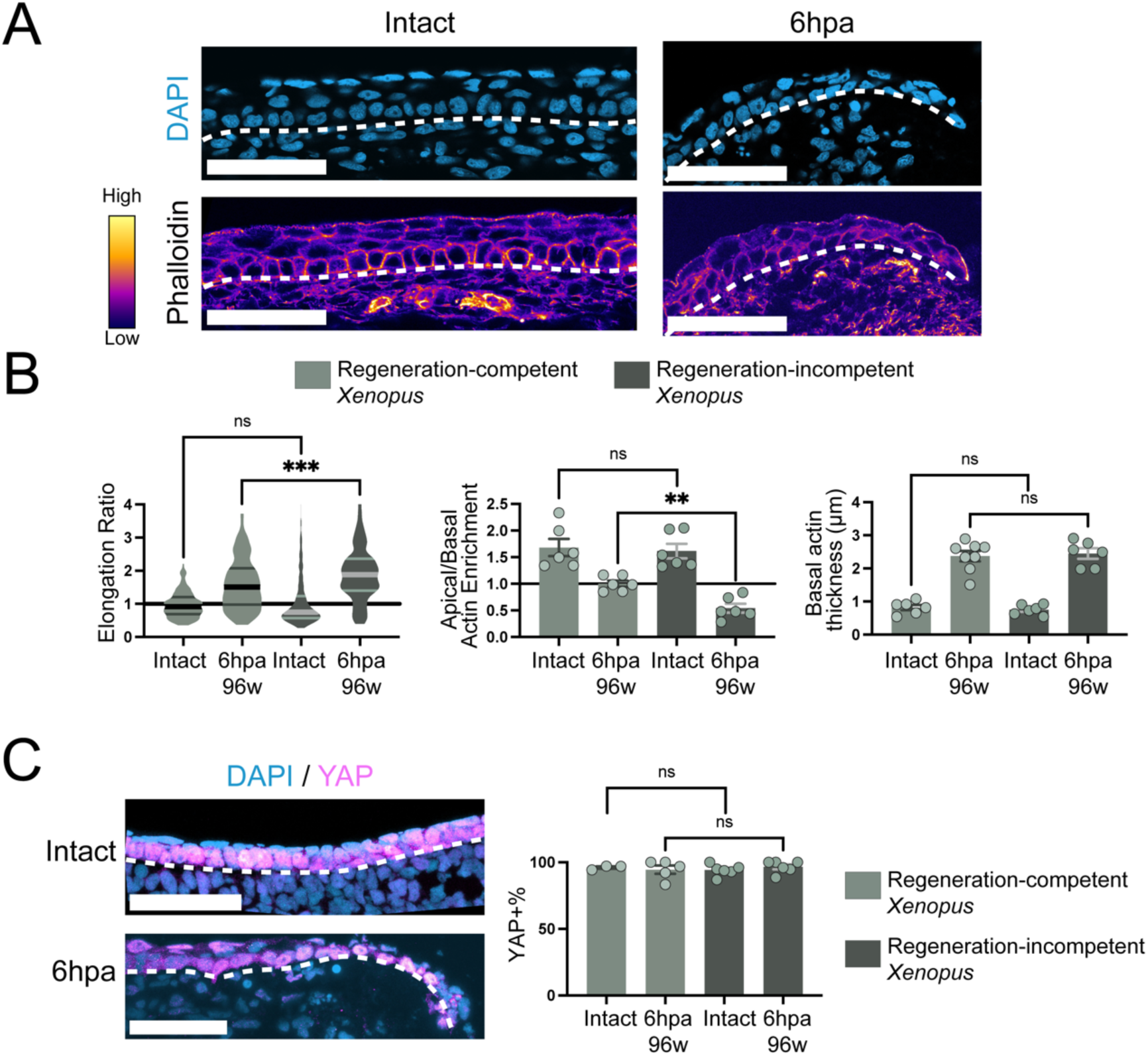
Regeneration-incompetent *Xenopus* tadpoles exhibit similar cell and actin reorganization and YAP activity as regeneration-competent tadpoles. **(A)** (Top) Confocal images of regeneration-incompetent *Xenopus* tadpole limbs, either intact or 6 hpa. Blue = DAPI. (Bottom) Phalloidin-stained confocal images of limbs under the same conditions. Scale bar = 50 μm. N = 3, total n = 9 for each condition. **(B)** (Left) Violin plot quantifying the elongation ratio of individual basal epithelial cells in *Xenopus* tadpoles under indicated conditions. (Middle) Bar plot showing apical-to-basal actin enrichment ratios in basal epithelial cells. Each dot represents individual sections. (Right) Bar plot of basal actin thickness in basal epithelial cells. Each dot represents individual sections. Note that regeneration-competent intact and 6 hpa samples are the same as in Figure 2A, B, F and are included here for comparison with regeneration-incompetent tadpoles. Regeneration-competent intact N=3, total n **=** 6, total quantified cell n = 119; regeneration-competent 6hpa N=3, n = 7, total quantified cell n = 139; regeneration-incompetent intact N=3, n = 6, total quantified cell n = 120; regeneration-incompetent 6hpa N=3, n = 6, total quantified cell n = 120. Total quantified cell number applies to elongation ratio. **(C)** Representative confocal images showing nuclear YAP immunofluorescence in *Xenopus* tadpoles under indicated conditions. Dashed lines outline the basement membrane. Scale bar = 50 μm. (Right) Bar plot quantifying distal basal epithelial cells with nuclear YAP. Note that intact regeneration-competent and 6 hpa sample data are the same as in Figure 2G and are included here for comparison with regeneration-incompetent tadpoles. Xenopus intact N = 3, n = 3; Xenopus 96w 6hpa N = 3, n=5; Intact regeneration-incompetent N=3, n=6; regeneration-incompetent 6hpa N=3, n=6.

**Figure S6.**
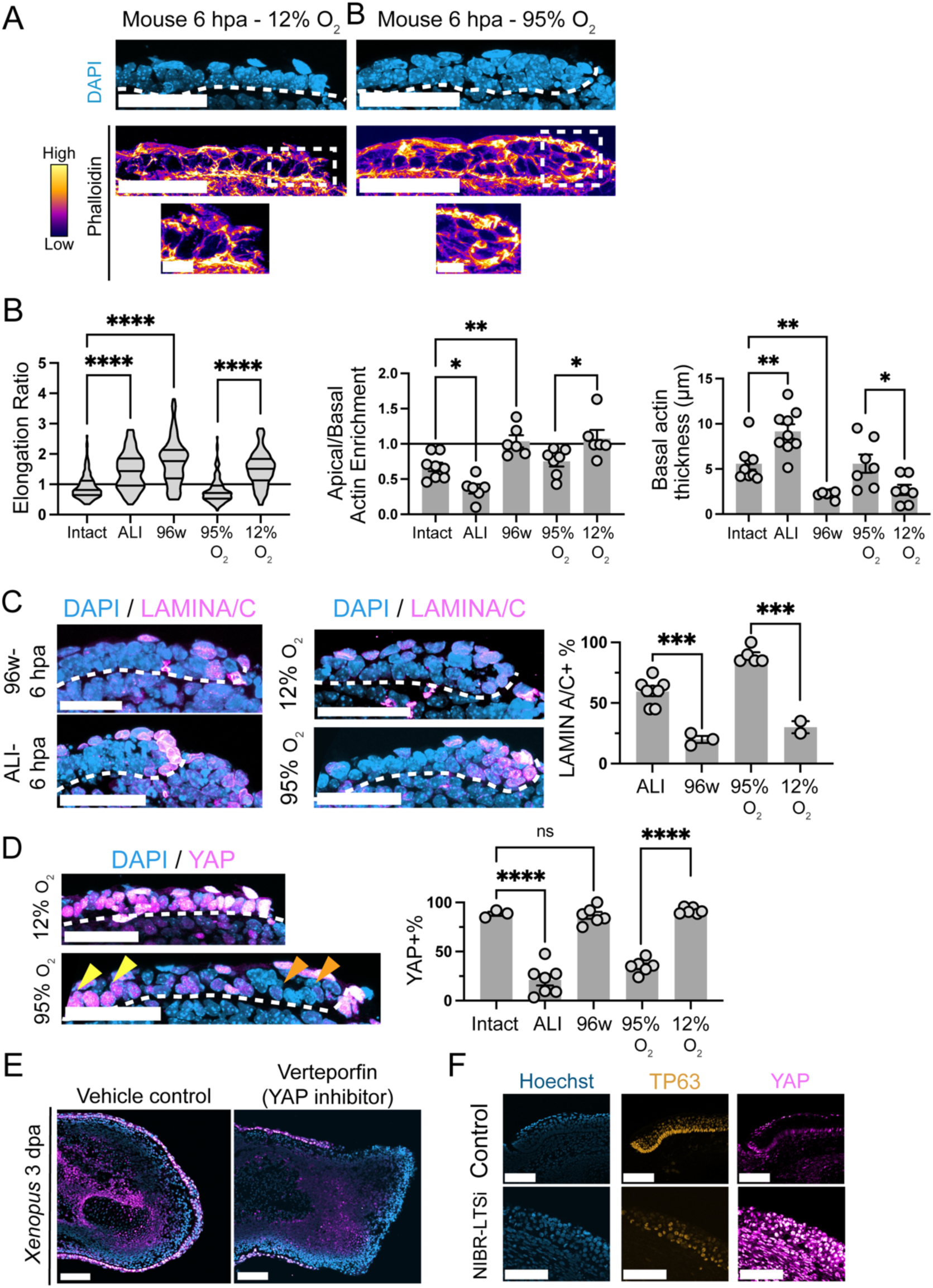
Low oxygen alters actin organization, reduces LAMIN/AC levels, and sustains active YAP in basal epithelial cells. **(A)** (Top) Confocal images of mouse limbs at 6 hpa under 12% or 95% oxygen culture conditions. Blue = DAPI. (Middle) Phalloidin-stained confocal images of mouse limbs under the same conditions. (Bottom) Zoomed regions (dashed lines) highlighting actin levels. N = 3, total n = 9. Scale bar = 50 μm; zoomed images = 10 μm. **(B)** (Left) Violin plot quantifying the elongation ratio of individual basal epithelial cells in mouse limbs under indicated conditions. Mouse intact N= 8 n= 159; Mouse ALI N= 6 n= 120; Mouse 6hpa N= 6 n= 119. (Middle) Bar plot of apical-to-basal actin enrichment ratios in basal epithelial cells. Each dot represents individual sections. N = 3, total n = 9. (Right) Bar plot of basal actin thickness in basal epithelial cells. Note that intact mouse, and ALI and 96w 6 hpa samples are the same as in Figure 2C-F and are included here for comparison with 12% oxygen and 95% oxygen levels. Each dot represents individual sections. N = 3, total n = 9 for both conditions. **(C)** (Left) Representative confocal images showing LAMINA/C immunofluorescence in mouse limbs under indicated conditions. Dashed lines outline the basement membrane. Scale bar = 50 μm. (Right) Bar plot quantifying basal epithelial cells with enriched LAMINA/C. Each dot represents individual sections. N = 3, total n = 9. **(D)** Representative confocal images showing nuclear YAP immunofluorescence in mouse limbs under indicated conditions. Dashed lines outline the basement membrane. Yellow arrows mark nuclear YAP in proximal cells, and orange arrows highlight its absence in distal basal epithelial cells. Scale bar = 50 μm. **(Right)** Bar plot quantifying distal basal epithelial cells with nuclear YAP. Each dot represents individual sections. Note that intact mouse ALI and 96w 6 hpa samples are the same as in Figure 2G and are included here for comparison with 12% oxygen and 95% oxygen levels. N = 3, total n = 9. **(E)** Representative 3 dpa *Xenopus* explant images treated with either vehicle control or Verteporfin. Note that the Verteporfin treatment image is the same as in Figure 2H and has been added here for comparison with the vehicle control. Blue: DAPI; Orange: TP63; Magenta: YAP. N = 3, total n = 9. Scale bar = 100 μm. **(F)** Representative confocal images of TP63-positive basal epithelial cells and YAP activity after NIBR-LTSi treatment in 3 dpa mouse explants cultured in ALI. Blue: DAPI; Orange: TP63; Magenta: YAP. N = 2, n = 6. Scale bar = 100 μm.

**Figure S7.**
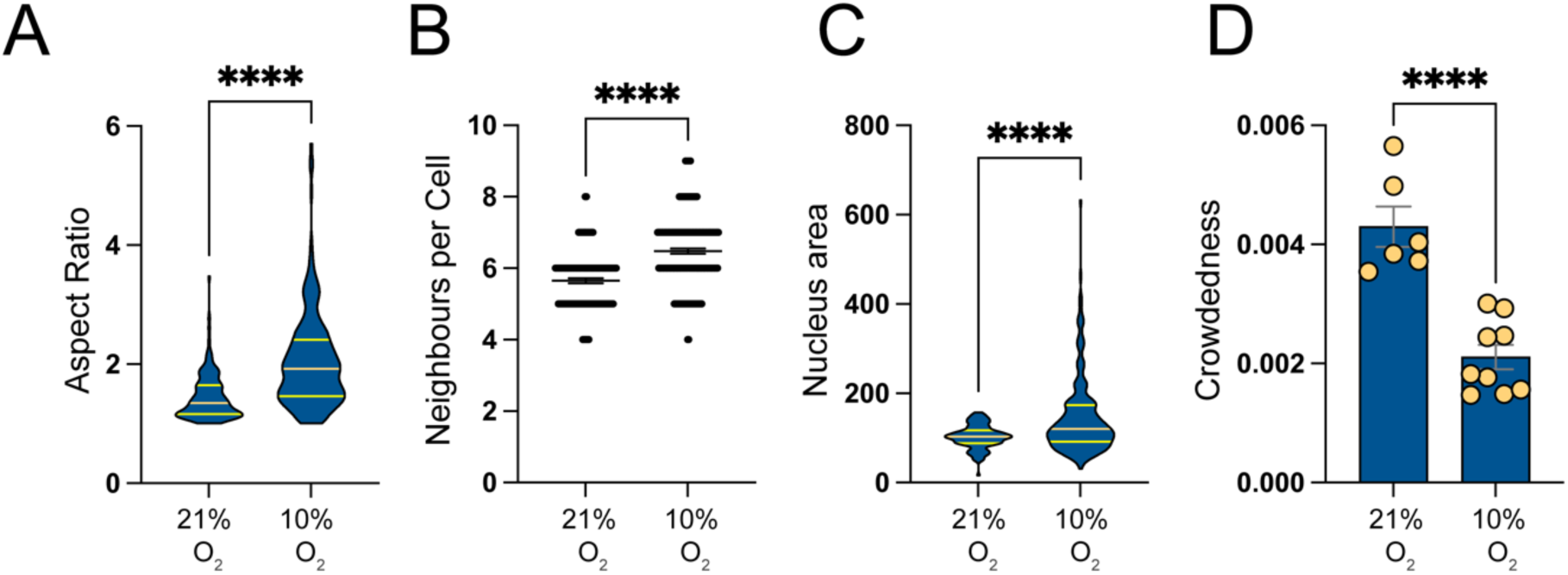
Quantifications for MDCK cell line cultured in 21% oxygen or 10% oxygen. **(A)** Violin plot quantifying the elongation ratio of MDCK cells in indicated conditions. 21% oxygen N=3, total wells. analyzed n=6, total quantified cell n= 120, 10% oxygen N=3, total wells analyzed n=8, total quantified cell n=180. **(B)** Scatter plot showing the neighbours per cells for MDCK cells in indicated conditions. 21% oxygen N=3, total wells. analyzed n=6, total quantified cell n= 120, 10% oxygen N=3, total wells analyzed n=8, total quantified cell n=180. **(C)** Violin plot quantifying the nucleus area of MDCK cells in indicated conditions. 21% oxygen N=3, total wells. analyzed n=6, total quantified cell n= 120, 10% oxygen N=3, total wells analyzed n=8, total quantified cell n=180. **(D)** Barplot showing cell numbers in µm² in indicated conditions. Each dot represents individual wells. 21% oxygen N=3, total wells. analyzed n=6, total quantified cell n= 120, 10% oxygen N=3, total wells analyzed n=8, total quantified cell n=180.

**Figure S8.**
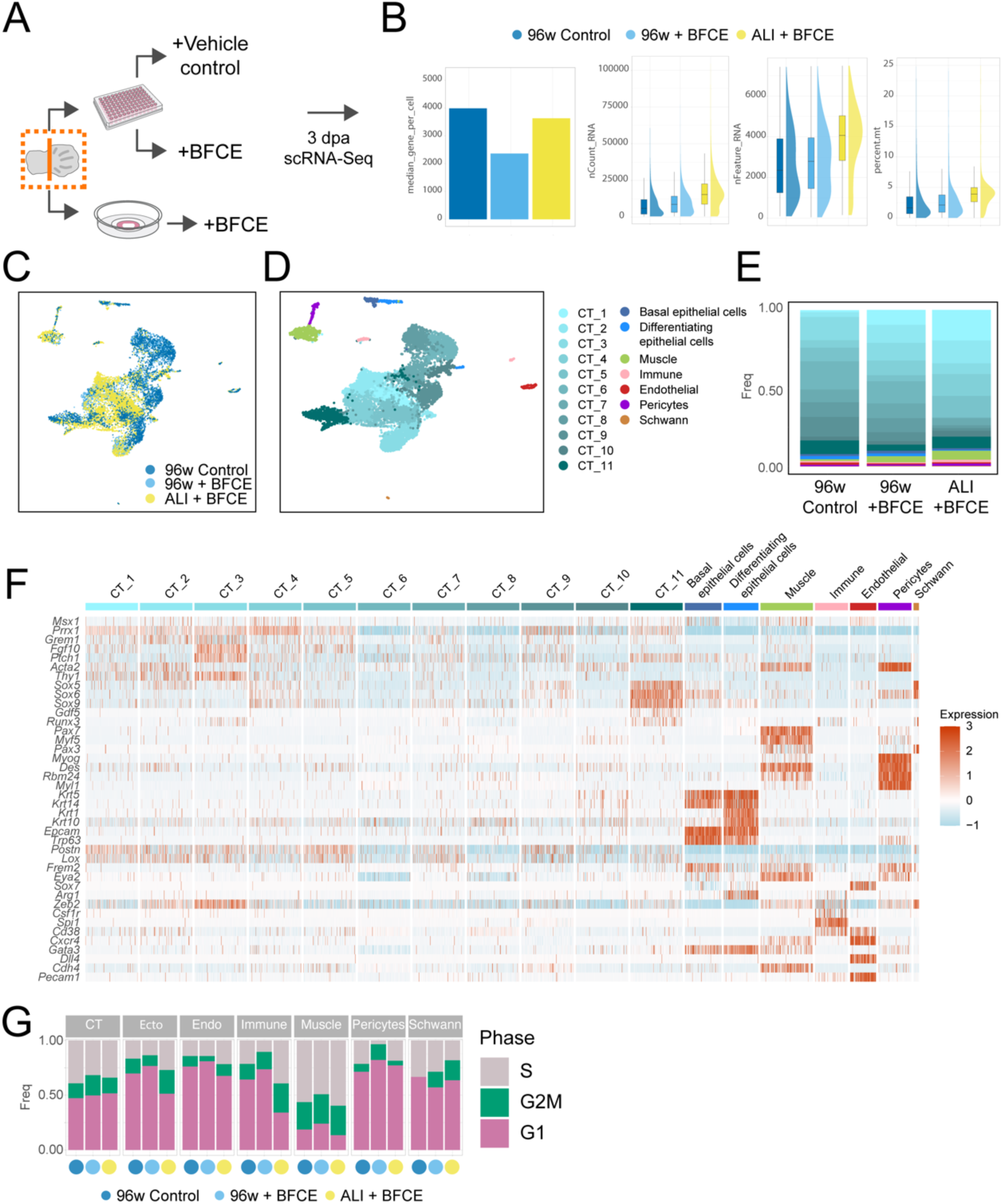
Experimental design, quality control, and additional details of 3 dpa scRNA-Seq experiments for 96w control, 96w with BFCE treatment, and ALI with BFCE treatment. **(A)** Schematic representation of the scRNA-Seq experimental design. BFCE: BMP4 + FGF10 + Chiron + EGF. **(B)** Quality control metrics for scRNA-Seq. Samples are color-coded as indicated above the figure. **(C)** Sample contributions to the integrated UMAP. Samples are color-coded as indicated in the figure. **(D)** Integrated UMAP representation of 3 dpa scRNA-Seq samples from (1) 96w with vehicle control, (2) 96w treated with BFCE, and (3) ALI treated with BFCE. Note that this panel is the same as Fig 3B but includes full annotation of cell clusters. **(E)** Bar plots showing the percentage of individual cell clusters in the scRNA-Seq data for each sample. Note that CT11 has high expression of cartilage markers (e.g. *Sox5, Sox6, Sox9*) and is labelled darker blue. **(F)** Heatmap depicting marker gene expression across different cell clusters. **(G)** Cell cycle scores derived from scRNA-Seq data, shown for each condition and grouped by major cell types. Ecto = basal epithelial cells. Endo = endothelial cells.

**Figure S9.**
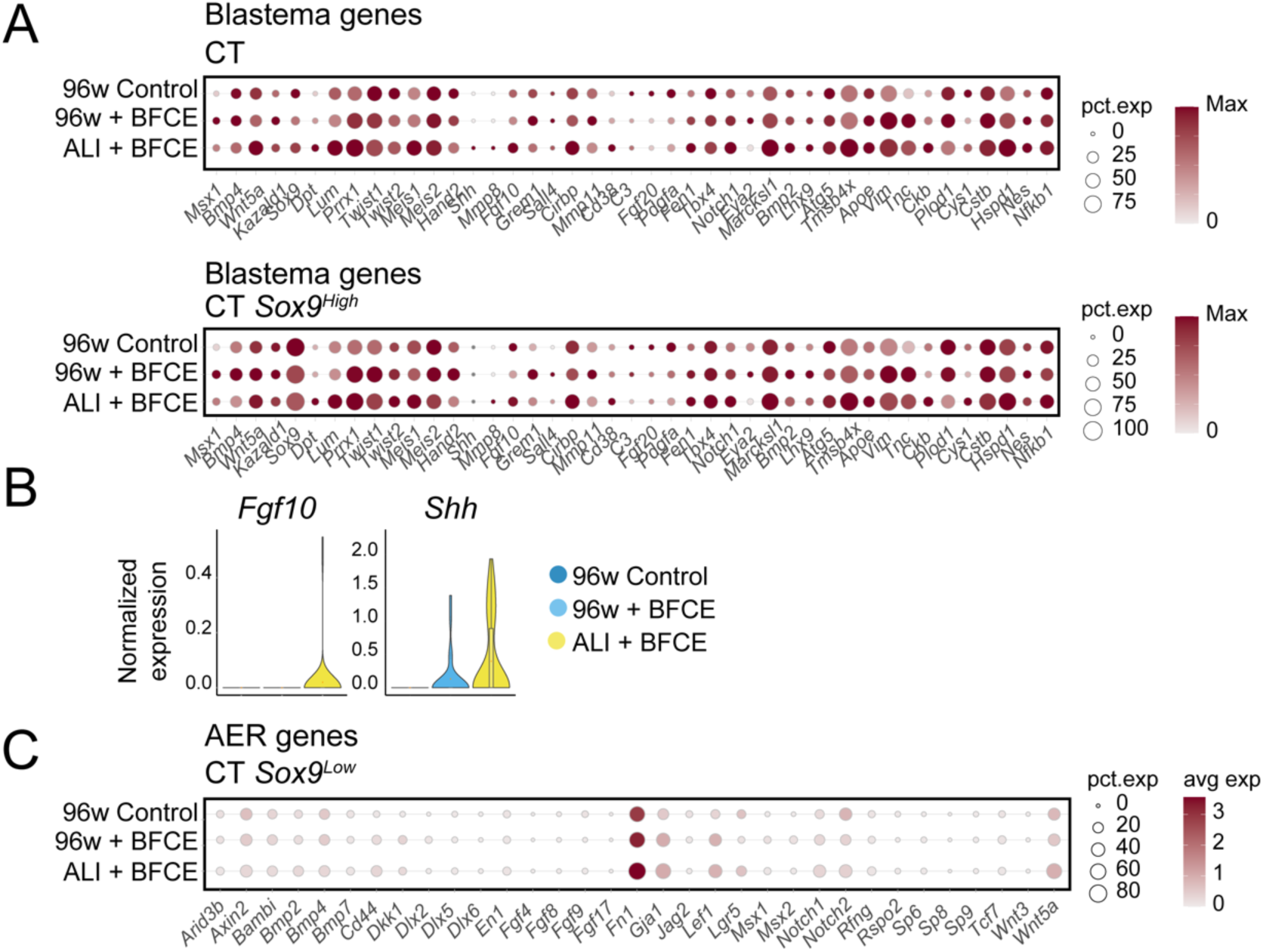
Blastema and AER gene expressions in different populations in scRNA-Seq data from Figure S8. **(A)** Dot plot showing blastema gene expression in (top) all connective tissue (CT) clusters and (bottom) *Sox9^High^* (cartilage) CT clusters associated with chondrogenic fate across 96w control, 96w BFCE treatment, and ALI BFCE treatment conditions. **(B)** Violin plots showing *Fgf10* and *Shh* expression in basal epithelial cells across the indicated experimental groups. **(C)** Dot plot showing AER gene expression in *Sox9^Low^* (non-cartilage) CT clusters across 96w control, 96w BFCE treatment, and ALI BFCE treatment conditions.

**Figure S10.**
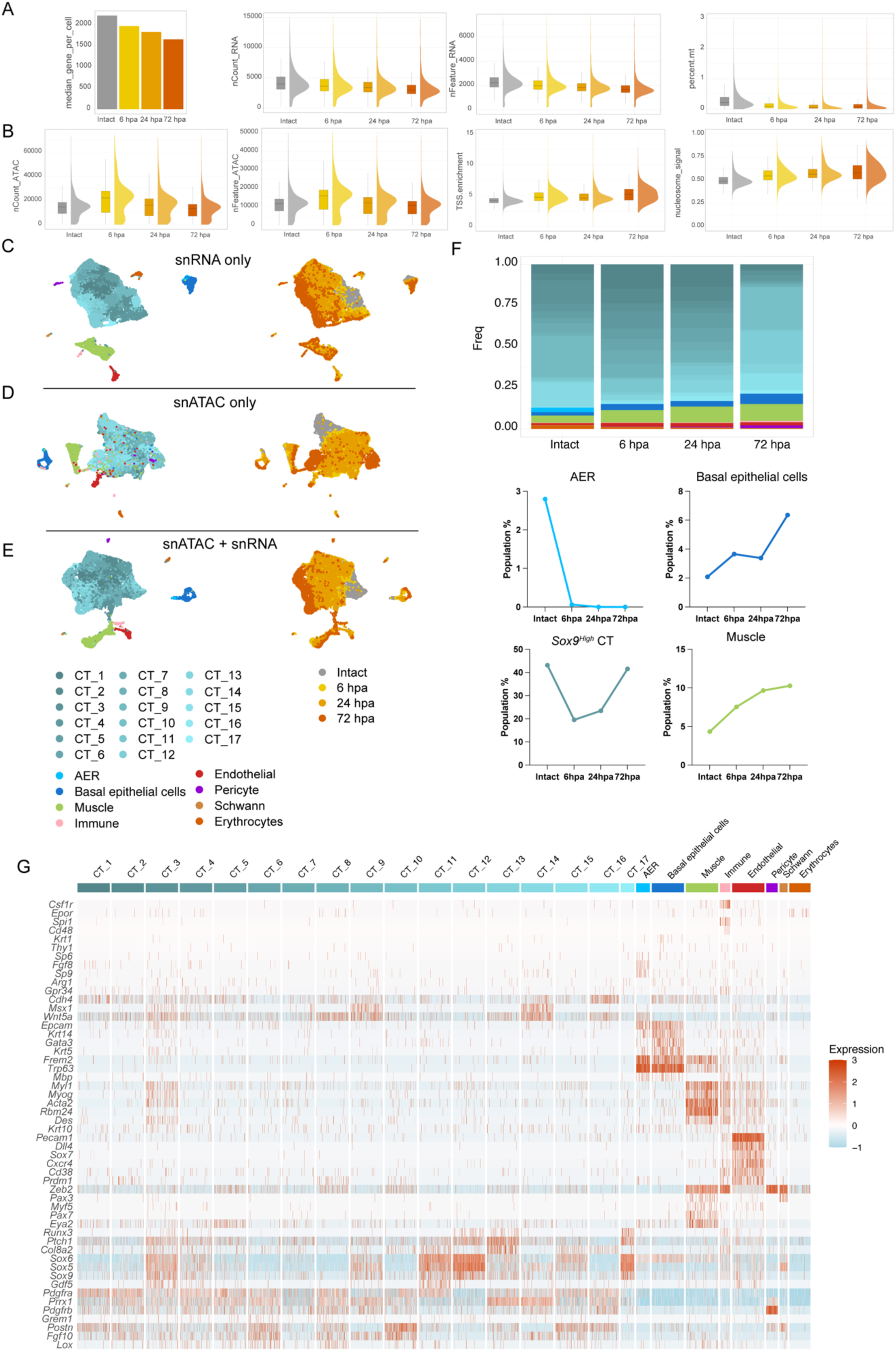
Quality control and additional details of scMultiome for the time-course ALI mouse explant experiment. **(A)** Quality control metrics for generated snRNA-Seq data. Samples are color-coded as gray: intact E12.5 limb; yellow: 6 hpa; orange: 24 hpa; red: 72 hpa. **(B)** Quality control metrics for generated snATAC-Seq data. Samples are color-coded as gray: intact E12.5 limb; yellow: 6 hpa; orange: 24 hpa; red: 72 hpa. **(C)** (Left) UMAP representation of snRNA-Seq-only data from the analyzed scMultiome experiment. Note that this panel is the same as Fig 4A but includes the full annotation of cell clusters at the bottom. (Right) Sample contributions to the integrated UMAP for snRNA-Seq, color-coded as gray: intact E12.5 limb; yellow: 6 hpa; orange: 24 hpa; red: 72 hpa. **(D)** (Left) UMAP representation of snATAC-Seq-only data from the analyzed scMultiome experiment. (Right) Sample contributions to the integrated UMAP for snRNA-Seq, color-coded as gray: intact E12.5 limb; yellow: 6 hpa; orange: 24 hpa; red: 72 hpa. **(E)** (Left) UMAP representation of combined snRNA and snATAC-Seq data from the analyzed scMultiome experiment. (Right) Sample contributions to the integrated UMAP for snRNA-Seq, color-coded as gray: intact E12.5 limb; yellow: 6 hpa; orange: 24 hpa; red: 72 hpa. **(F)** (Top) Percentage of indicated cell clusters in snRNA-Seq data for each sample. (Bottom) Line plots showing the population percentages of relevant cell populations across analyzed time points*.Sox9^High^*populations are clusters 3, 9, 11, 12, 13, 15, and 17. **(G)** Heatmap showing marker gene expression across different clusters.

**Figure S11.**
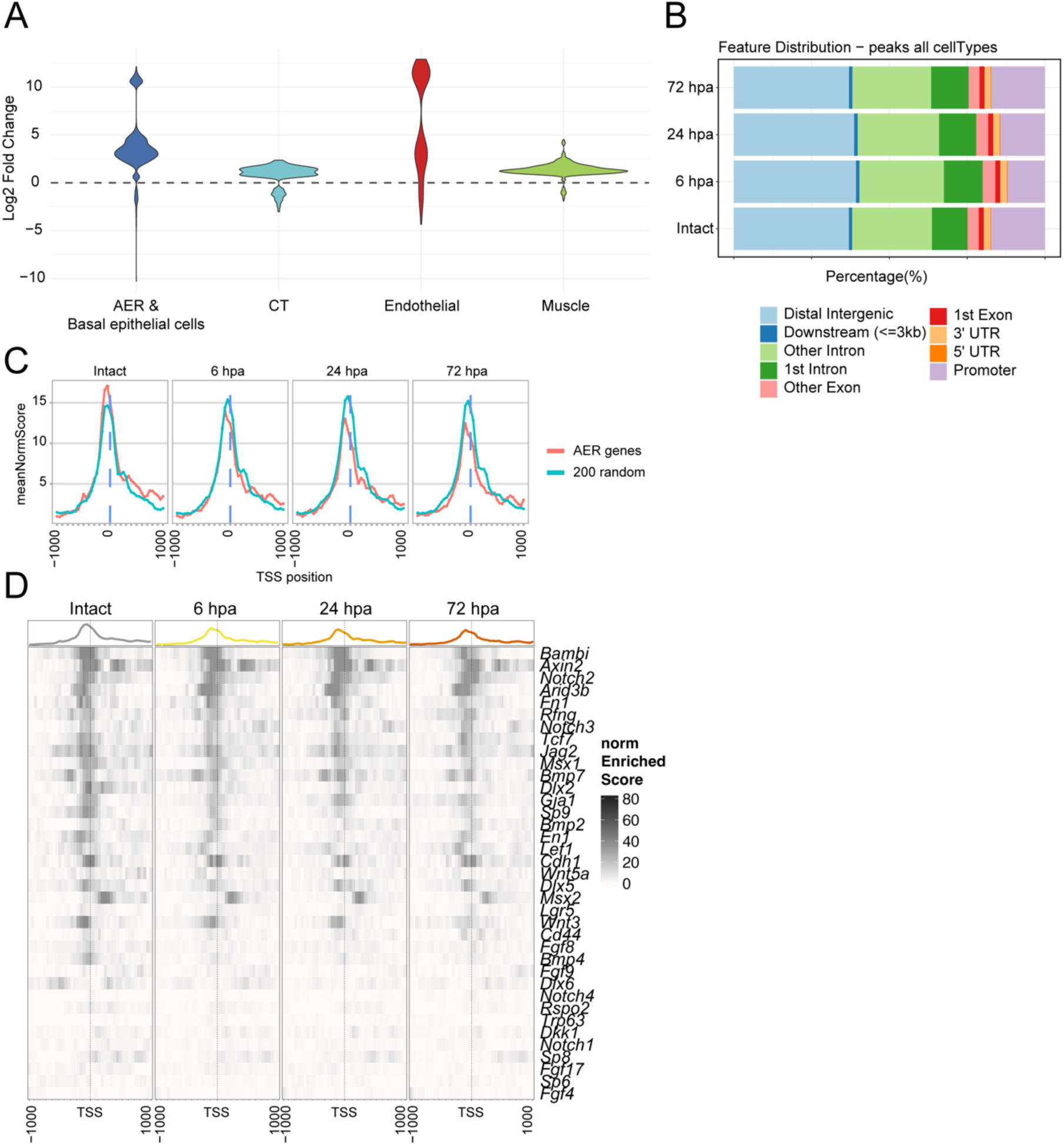
Chromatin accessibility changes in the scMultiome experiment from Figure S10. **(A)** Violin plots showing the differential chromatin accessibility between intact and 6 hpa samples for major cell types. **(B)** Percentage barplot showing the distribution of open chromatin regions across genomic features and for all datasets. **(C)** Line plot illustrating the enrichment profiles for the AER gene set and 200 random genes across time points. Note that this data is the same as Fig 4B-C but is overlaid here to emphasize relative changes at each time point. **(D)** Heatmap depicting AER gene set chromatin accessibility at each time point in basal epithelial cells.

**Figure S12.**
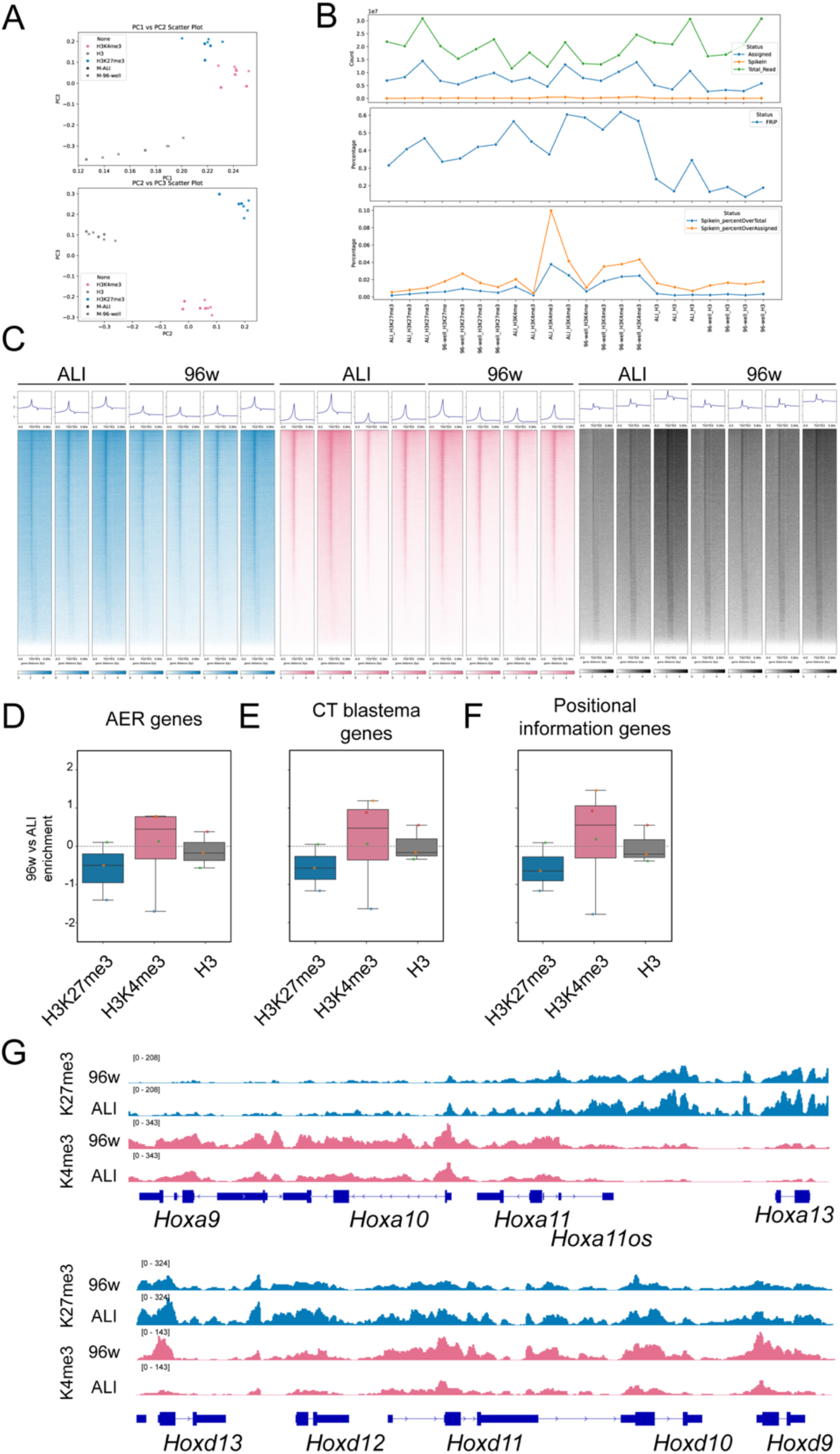
Quality control and additional details of CUT&Tag experiments for 1 dpa ALI and 96w mouse explant experiments. **(A)** Principal component analysis (PCA) of processed CUT&Tag samples. **(B)** Quality control metrics for CUT&Tag experiments, including assigned reads and spike-in levels. Note that one ALI H3K4me3 sample shows an overrepresentation of spike-ins. **(C)** Heatmaps of TSS peaks across individual replicates. **(D-F)** Box plots comparing the enrichment ratios for H3K4me3, H3K27me3, and total H3 modifications between 96w plates and ALI culture conditions for AER, blastema, and positional information gene sets. **(G)** Example genome browser snapshots of representative replicates showing enrichment patterns for *HoxA* and *HoxD* clusters in mouse explants.

**Figure S13.**
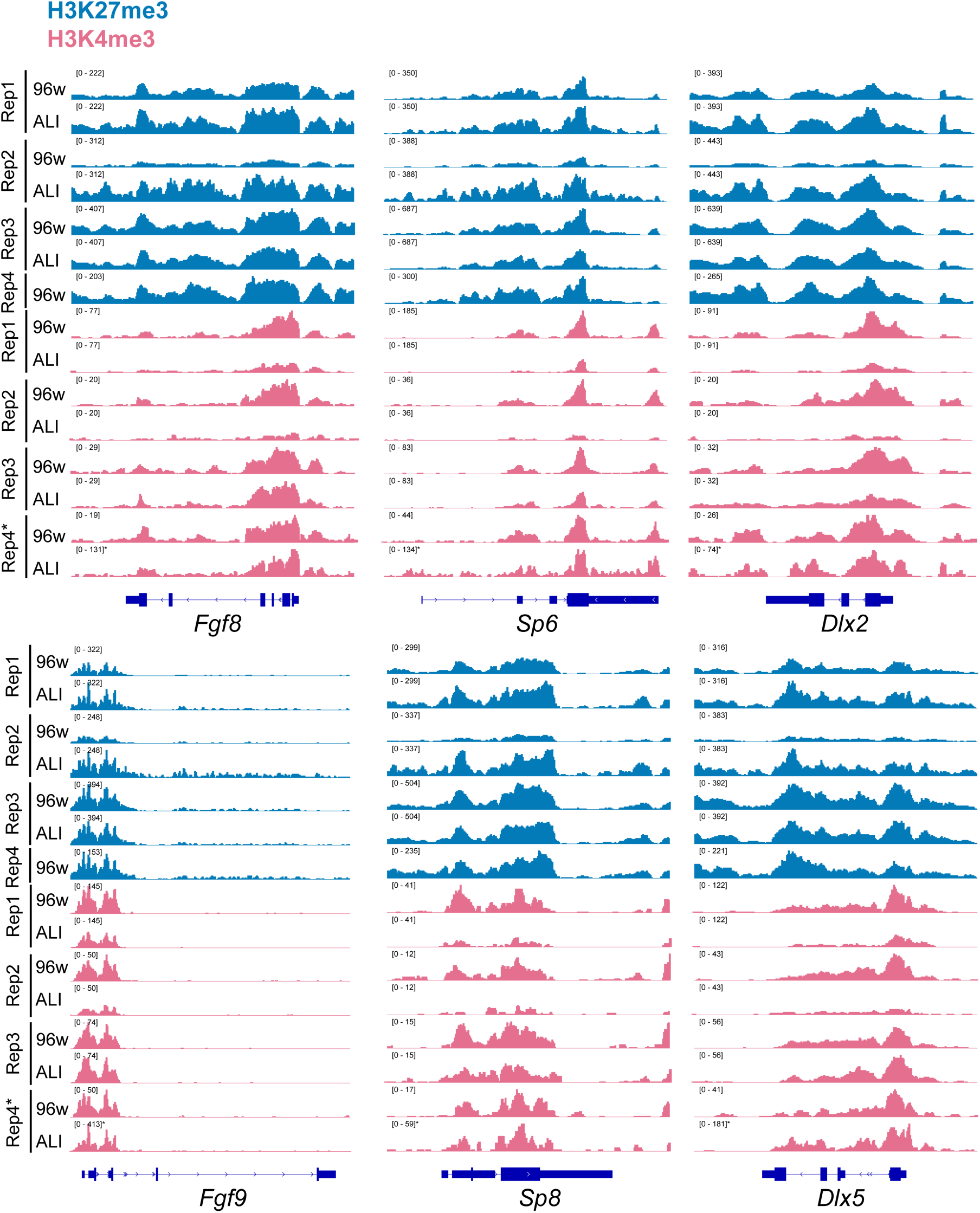
Genome browser track snapshots for each replicate of example AER genes in mouse 96w and ALI 1 dpa CUT&Tag samples. Genome browser tracks showing AER gene enrichment profiles across all replicates. Note that all replicates were scaled with their pairing condition, except for two samples: (1) Replicate 4 (Rep4) of H3K27me3 had only 96w samples and was autoscaled, and (2) Rep4 of the H3K4me3 ALI condition exhibited a significantly increased range. Thus, autoscaling is used for that group.

**Figure S14.**
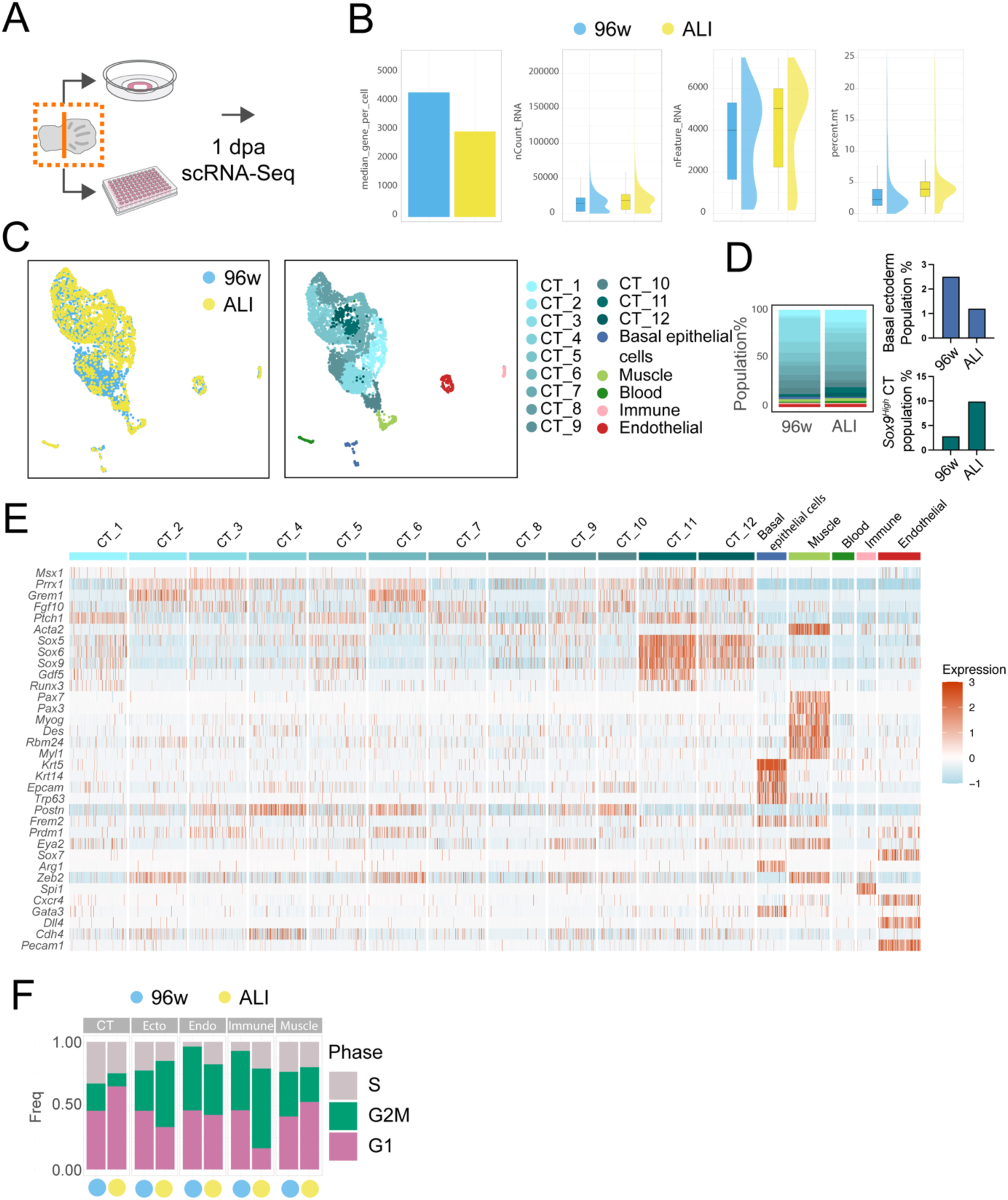
Experimental design, quality control, and additional details of scRNA-Seq experiments for 1 dpa ALI and 96w mouse explant experiments. **(A)** Schematic representation of the scRNA-Seq experimental design. **(B)** Quality control metrics for generated scRNA-Seq data. Samples are color-coded as indicated in the figure. **(C)** Sample contributions to the integrated UMAP, with blue representing 96w and yellow representing ALI conditions. **(D)** (Left) Integrated UMAP representation of 3 dpa scRNA-Seq samples from (1) 96w with vehicle control, (2) 96w treated with BFCE, and (3) ALI treated with BFCE. *Sox9^High^* populations are clusters 11 and 12. Note that this panel is the same as Fig 4H but includes the full annotation of cell clusters (right). **(E)** Heatmap showing marker gene expression across different clusters. **(F)** Cell cycle scores derived from scRNA-Seq data, divided by condition and shown for groups of clusters. Ecto = basal epithelial cells. Endo = endothelial cells.

**Figure S15.**
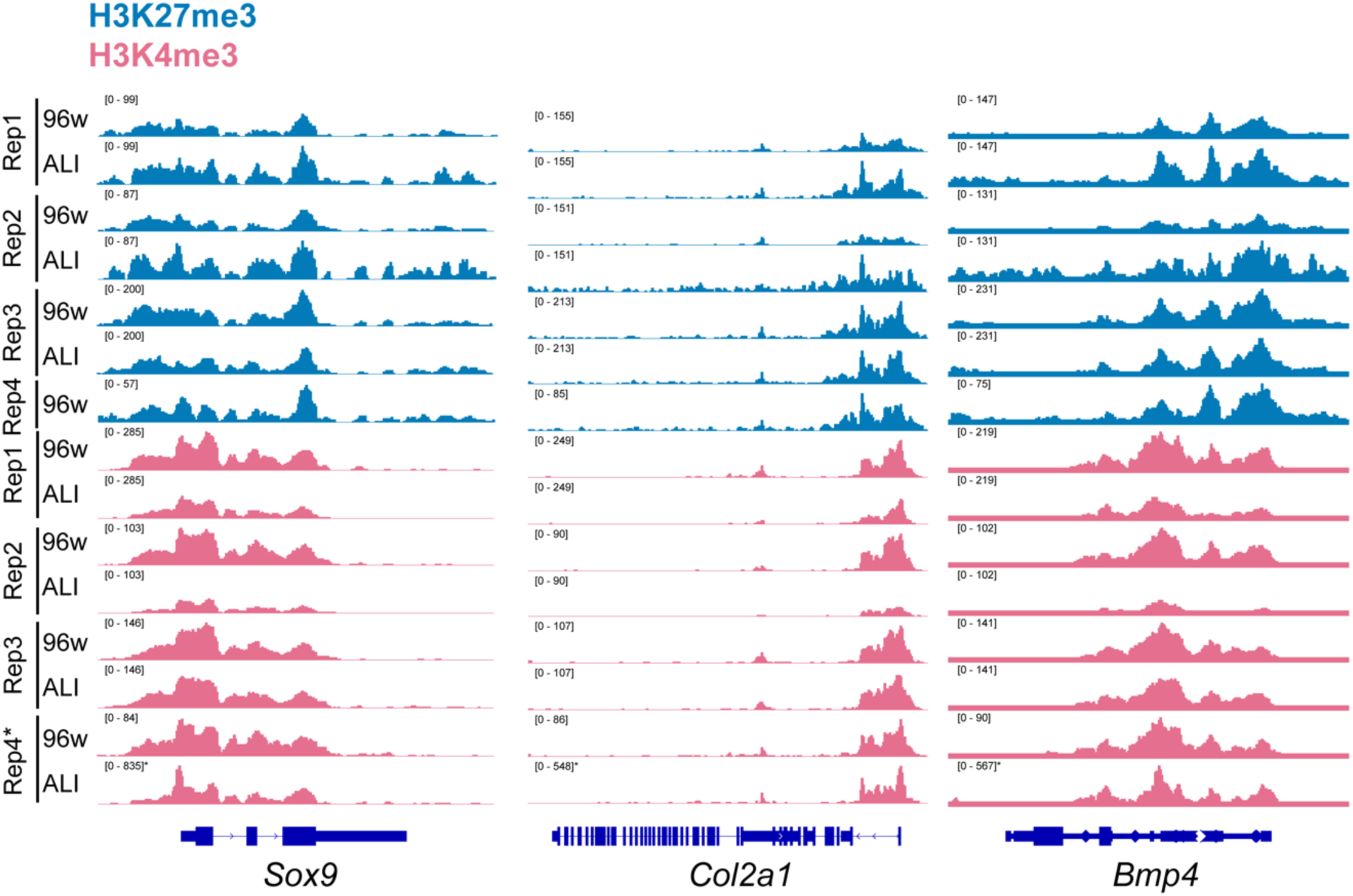
Genome browser track snapshots for each replicate of example chondrogenic genes in mouse 96w and ALI 1 dpa CUT&Tag samples. Genome browser tracks showing cartilage formation-related gene enrichment profiles across replicates. Note that all replicates were scaled with their pairing condition, except for two samples: (1) Replicate 4 (Rep4) of H3K27me3 had only 96w samples and was autoscaled, and (2) Rep4 of the H3K4me3 ALI condition exhibited a significantly increased range. Thus, autoscaling is used for that group.

**Figure S16.**
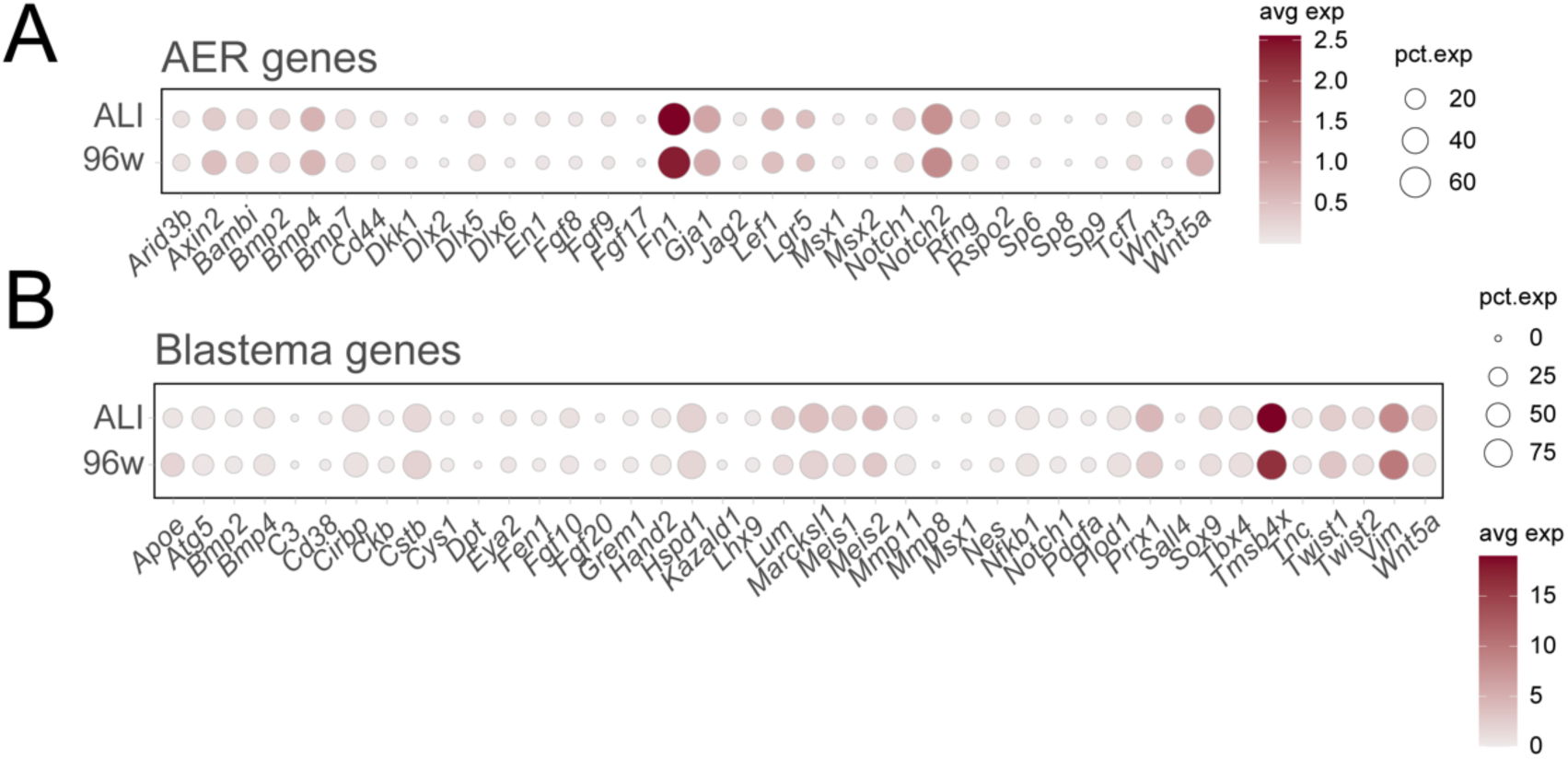
AER and blastema gene expressions in mouse ALI and 96w 1 dpa scRNA-Seq samples. **(A)** Dot plot illustrating AER gene expression in 1 dpa ALI and 96w mouse explants. **(B)** Dot plot illustrating blastema gene expression in 1 dpa ALI and 96w mouse explants.

**Figure S17.**
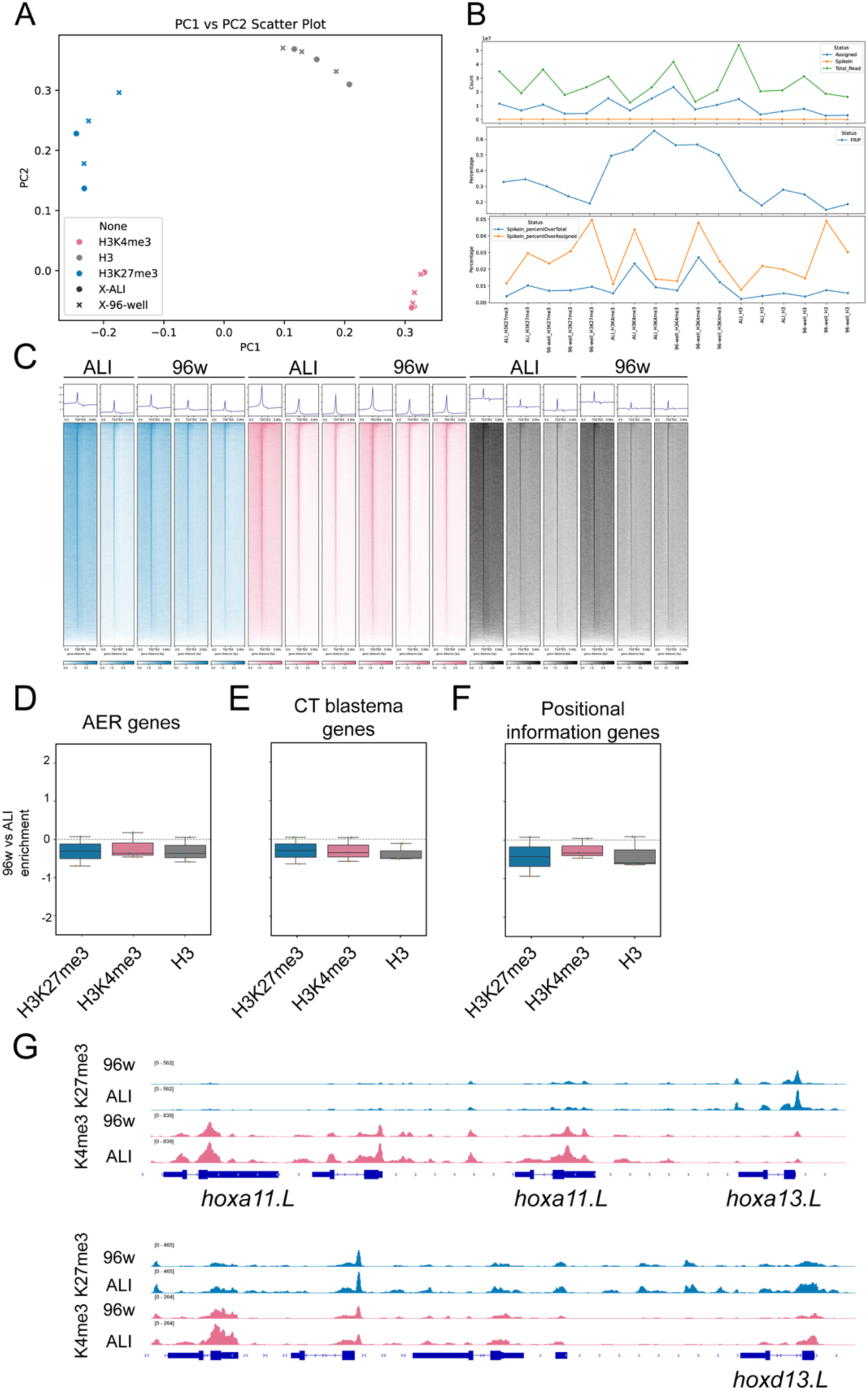
Quality control and additional details of CUT&Tag experiments for 1 dpa ALI and 96w *Xenopus* explant experiments. **(A)** Principal component analysis (PCA) of processed CUT&Tag samples. **(B)** Quality control metrics for CUT&Tag experiments, including assigned reads and spike-in levels. **(C)** Heatmaps of TSS peaks across individual replicates. **(D-F)** Box plots comparing the enrichment ratios for H3K4me3, H3K27me3, and total H3 modifications between 96w and ALI culture conditions for AER, blastema, and positional information gene sets. **(G)** Example genome browser snapshots of a representative replicates for *HoxA* and *HoxD* clusters in *Xenopus* explants.

**Figure S18.**
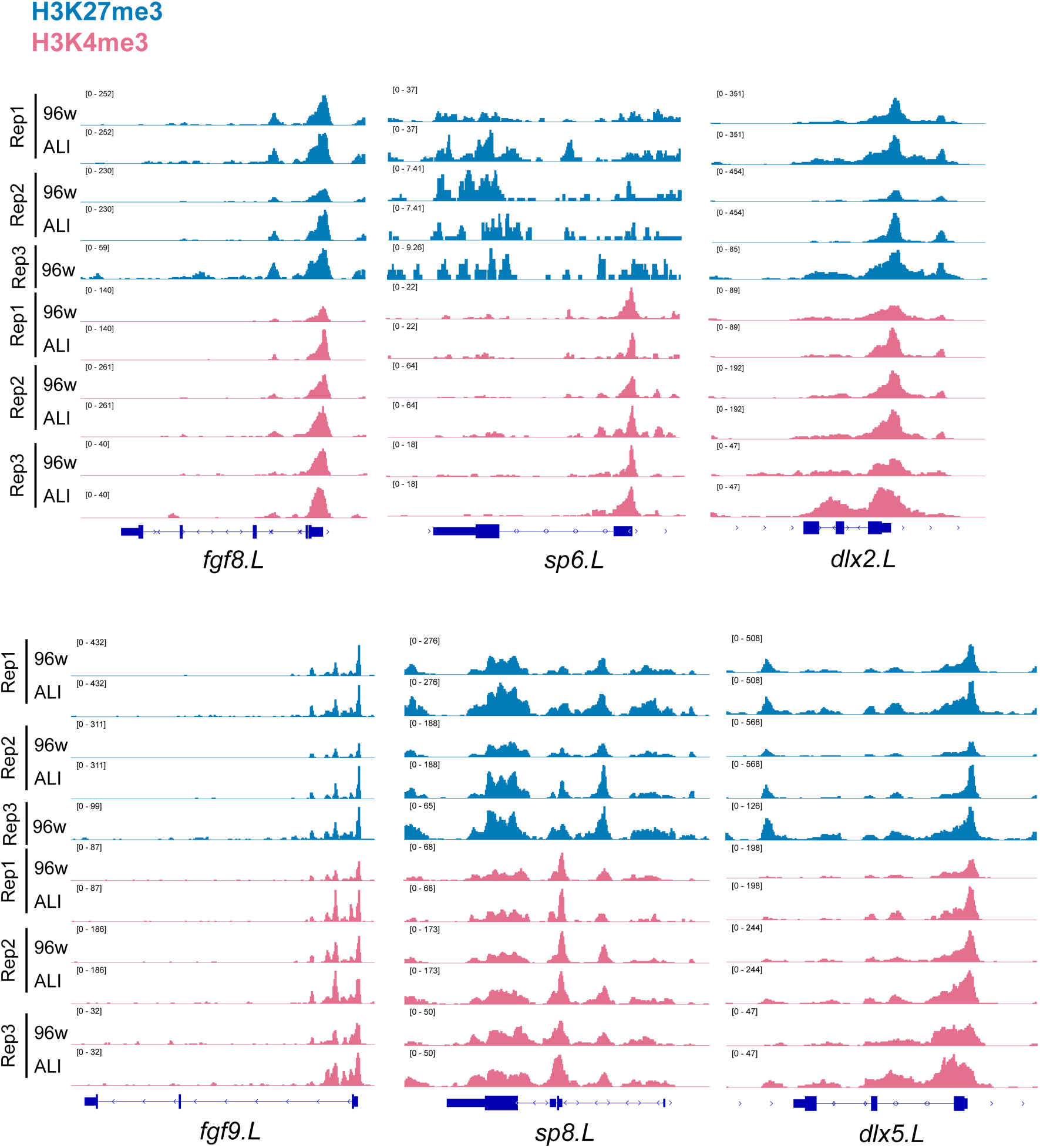
Genome browser track snapshots for each replicate of example AER genes in *Xenopus* 96w and ALI 1 dpa CUT&Tag samples. Genome browser tracks showing AER gene enrichment profiles across all replicates. Note that all replicates were scaled with their pairing condition, except for Rep3 of H3K27me3, which had only 96w samples and was autoscaled.

**Figure S19.**
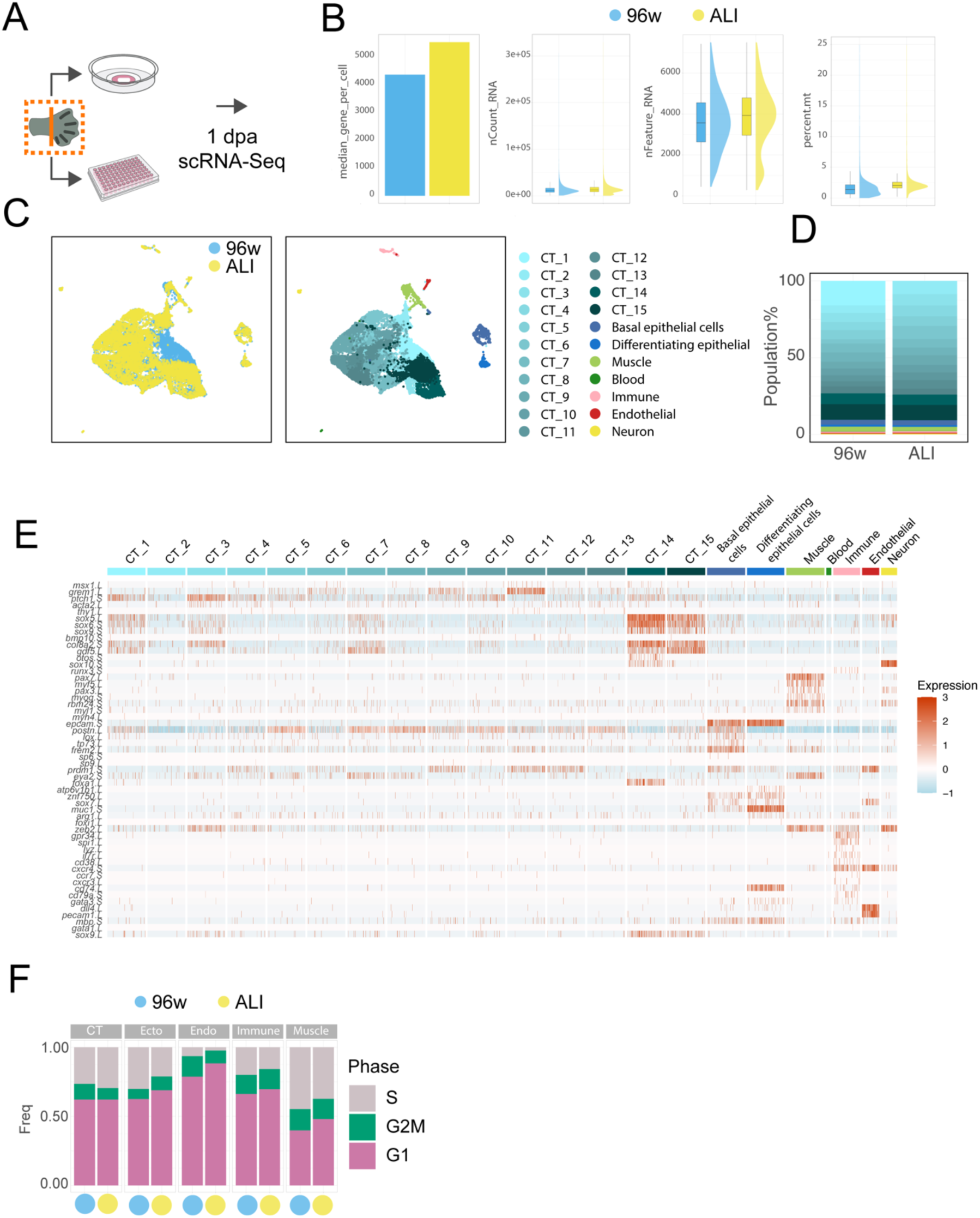
Experimental design, quality control, and additional details of scRNA-Seq experiments for 1 dpa ALI and 96w plate *Xenopus* explant experiments. **(A)** Schematic representation of the scRNA-Seq experimental design. **(B)** Quality control metrics for generated scRNA-Seq data. Samples are color-coded as indicated above the figure. **(C)** (Left) Sample contributions to the integrated UMAP. Samples are color-coded as indicated in the figure. (Right) Integrated UMAP representation of 1 dpa scRNA-Seq samples. Note that this figure is the same as Fig 5J but includes full annotation of cell clusters. **(E)** Heatmap showing marker gene expression across different clusters. **(F)** Cell cycle scores derived from scRNA-Seq data, divided by condition and shown for groups of clusters. Ecto = basal epithelial cells. Endo = endothelial cells.

**Figure S20.**
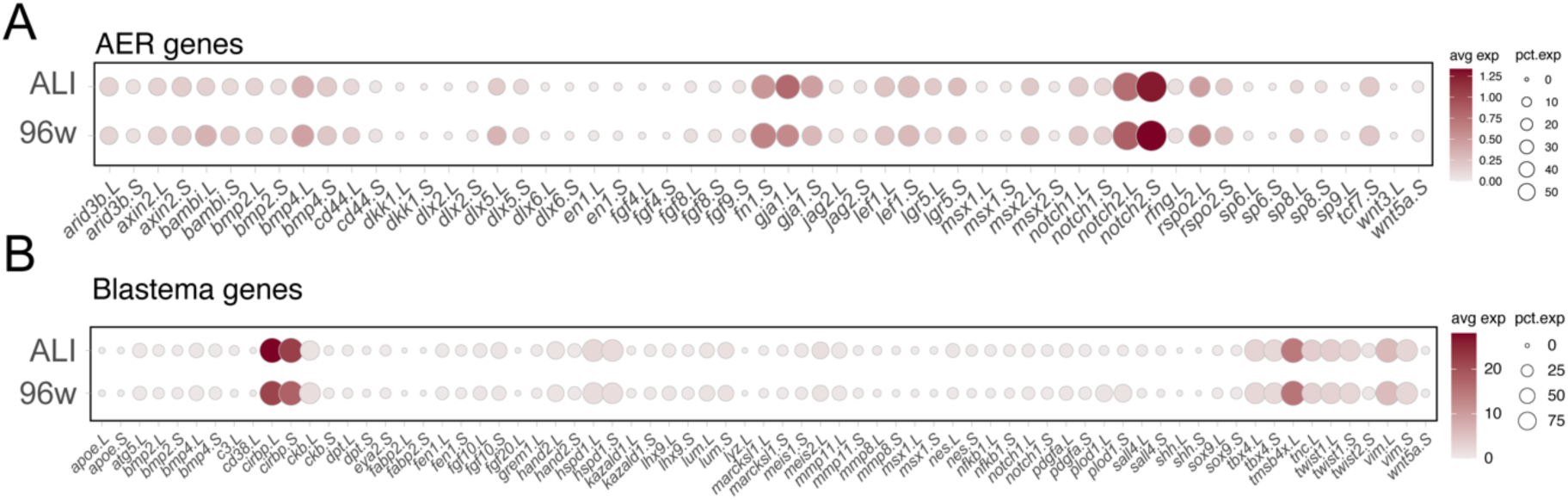
AER and blastema gene expressions in *Xenopus* ALI and 96w 1 dpa scRNA-Seq samples. **(A)** Dot plot illustrating AER gene expression in 1 dpa ALI and 96w *Xenopus* explants. **(B)** Dot plot illustrating blastema gene expression in 1 dpa ALI and 96w *Xenopus* explants.

**Figure S21.**
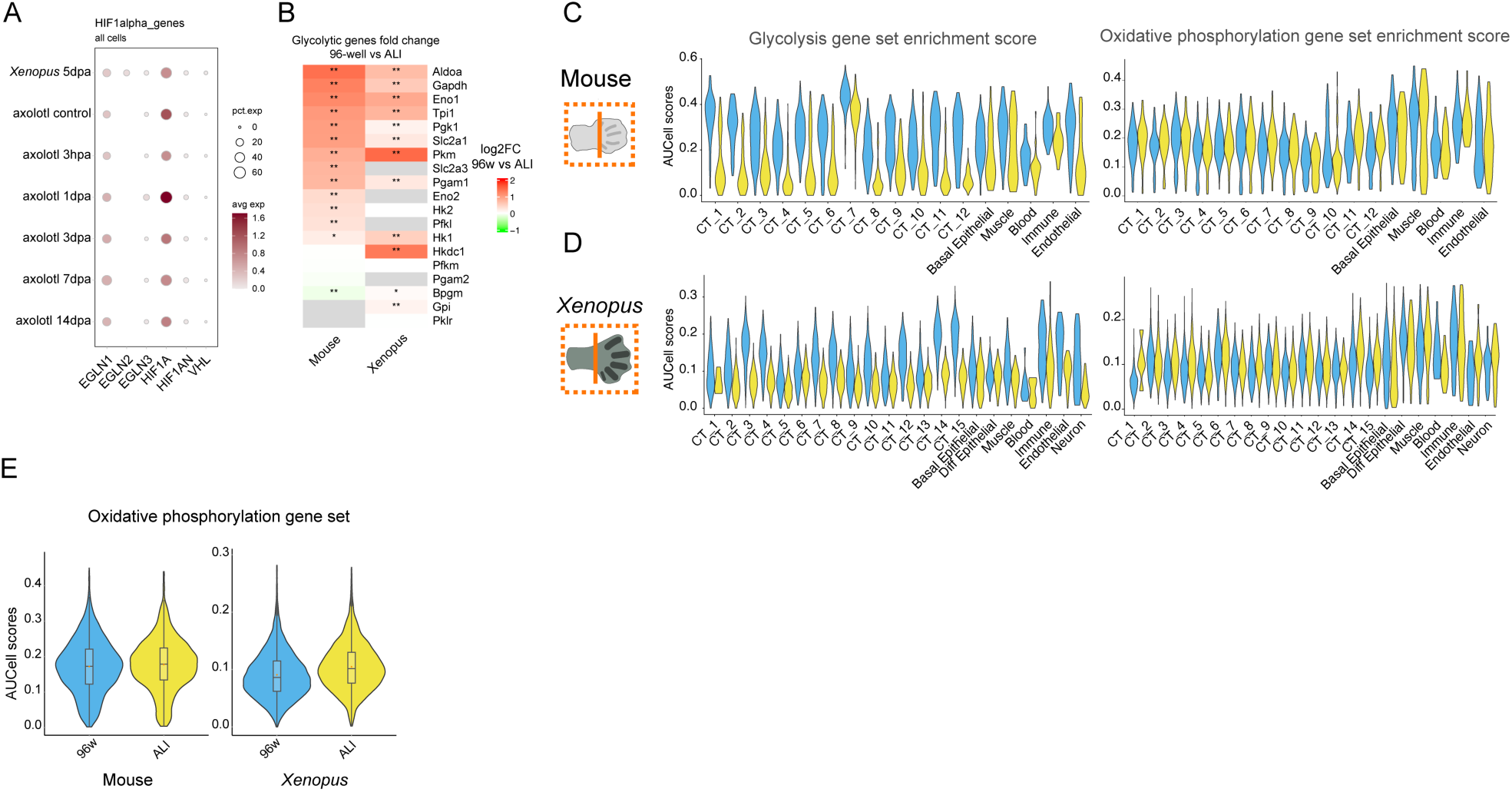
HIF1A regulator expression patterns during amphibian limb regeneration and glycolysis versus oxidative phosphorylation gene set expressions. **(A)** Dot plot comparing HIF1A pathway gene expression profiles across publicly available *Xenopus* and axolotl limb regeneration scRNA-seq datasets. **(B)** Heatmap showing fold changes in glycolysis gene expression between ALI and 96w for mouse and *Xenopus* 1 dpa scRNA-seq datasets. * in this figure denotes adjusted p-value < 0.05, and ** in this figure denotes adjusted p value <0.001. **(C)** Violin plots showing glycolysis gene set expression in 1 dpa scRNA-seq datasets across individual cell clusters for mouse explants in ALI and 96w. **(D)** Violin plots showing glycolysis gene set expression in 1 dpa scRNA-seq datasets across individual cell clusters for *Xenopus* explants in ALI and 96w. **(E)** Violin plots illustrating oxidative phosphorylation gene set expression in 1 dpa scRNA-seq datasets for *Xenopus* and mouse explants in ALI and 96w.

